# Gene Gradients Reveal Directed Structural Connectivity Across Species

**DOI:** 10.64898/2026.05.05.723068

**Authors:** Benjamin S Sipes, Fahimeh Arab, Srikantan S Nagarajan, Ashish Raj

## Abstract

Studying the brain’s inter-regional structural connectivity (SC) has transformed neuroscience [12], yet noninvasive neuroimaging remains blind to the directionality of neural wiring. Structural directionality must influence observed brain dynamics, but estimating edge-level directionality from brain function is ill-posed and underdetermined [31]. Here, we show that gene co-expression gradients define a low-dimensional biological manifold that locally biases brain regions to act as network sources or sinks. Embedding gene gradients into a linear, higher-order network diffusion model defines a gene-structure-function relationship that recovers ground-truth directionality in *C. elegans* (*r* = 0.70), mouse (*r* = 0.57), and macaque (*r* = 0.46). Applying this framework to 770 healthy young adults from the Human Connectome Project yields subject-specific directed connectomes that reproduce established corticothalamic asymmetries and reveal distinct gene ontologies associated with network sources and sinks. Furthermore, integrating directed structure with observed functional covariance enables us to derive a principled measure of directed functional connectivity, termed “angular flow” (AF), related to probability angular momentum in non-equilibrium statistical mechanics. Regional AF recapitulates the principal unimodal–multimodal gradient of functional connectivity, suggesting that this macroscale functional hierarchy emerges from a source–sink organization of signal flow through the brain. This framework resolves a long-standing limitation in connectomics and opens new opportunities to study structural directionality and non-equilibrium brain dynamics in health and disease.

## Introduction

Mapping whole-brain structural connectivity (SC) is a central goal in neuroscience. The first completely mapped “connectome” was in the nematode, *Caenorhabditis elegans*, comprising 302 neurons [101]. Critically, this connectome is *directional*—it establishes which neurons transmit action potentials to downstream neurons. Directionality is also fundamental to other aspects of neural physiology; for example, different metabolic cargo and trophic factors undergo anterograde or retrograde axonal transport [55]. Recent work suggests that these transport processes shape the directional spread of pathological proteins, including tau in Alzheimer’s disease [90, 89]. Highly detailed directed connectomes have enabled computational models that provide deep mechanistic insights into brain processing and support large-scale *in silico* simulation of brains and behavior [77]. In mammals, however, whole-brain single-cell wiring diagrams remain inaccessible. Instead, invasive tract-tracing studies have been used to map directed connectivity between brain regions in mice and non-human primates [63, 9], but these techniques are not suited for mapping human brain connectivity. As a result, present efforts to map human SC rely largely on diffusion magnetic resonance imaging (dMRI) tractography, which can measure the strength of connections between regions, but not the directionality. Therefore, neuroscience is in need of a framework to estimate structural directionality by other means.

Prior work has found a biological link between genes and SC, suggesting that transcriptional expression may aid in estimating directionality. Molecular gradients are well known to guide axon development [104], and regional gene-expression patterns are systematically related to macroscale brain connectivity [6, 35]. Early work in rodents found that brain regions with similar gene expression profiles tend to have similar connectivity profiles, and that connected regions tend to exhibit greater co-expression than unconnected regions [26]. Gene expression also relates to structural hub organization: studies in *C. elegans* and mouse have identified distinctive transcriptional signatures of hub connectivity and elevated gene expression similarity between hubs [5, 30]. Similar relationships have been observed in humans, where regional gene expression and inter-region coexpression are significantly associated with both structural and functional connectivity [35, 39, 96]. Gene-expression topographies also mirror large-scale cortical network organization, consistent with a role in long-range corticocortical connectivity [50]. Importantly, studies in mouse have identified relationships between gene-expression and cell-type distributions and the direction of source-target axonal connectivity [20, 85]. Together, these findings suggest that gene co-expression maps may also contain latent information about directed SC.

Network diffusion models over the SC have also shown potential to reveal hints of latent directionality. Asymmetric communication emerges when signaling is modeled as a diffusive process on the network [60, 70]. Extending this idea to the whole-brain, decentralized communication measures including diffusion efficiency were used to infer group-level sender–receiver asymmetries from undirected structural connectomes [73]. More generally, network diffusion models relate correlations between brain signals to passive signal propagation through SC [1, 2, 24, 31]. Simulated asymmetric, afferent-normalized diffusion over structural networks has been shown to better constrain effective connectivity than conventional anatomical connectivity alone [82]. Similarly, noise-diffusion models often leverage the *Lyapunov Equation*, which describes the stationary covariance of a linear stochastic differential equation and has been used to infer directed effective connectivity from time-shifted functional covariance [31]. However, these approaches relied on group-level graph theoretical measures or aimed only to predict effective functional, not structural, connectivity. To our knowledge, there is no existing model that can estimate subject-specific structural directionality from *in vivo* neuroimaging.

Estimating directionality for each edge in a network with *n* nodes conventionally would require *n*^2^ parameters, making the problem ill-posed and underdetermined [31]. However, embedding transcriptional expression within a network diffusion model could provide a parsimonious and biologically principled parameterization for estimating the brain’s directed SC. Here, we introduce such a gene-structure-function model based on two hypotheses: 1) gene co-expression patterns can provide a parsimonious parameterization for directionality, and 2) there exists latent directional information contained in the undirected covariance of brain signals. Our model assumes that local gene expression biases regions toward being sources or sinks, and the relative bias between regions modulates SC. We learn gene expression weights using a cost function based on the Lyapunov Equation, which fits the model under the assumption that empirical zero-lag functional connectivity is stationary. The result of our model fitting provides a prediction for the directed SC.

Our results establish a biologically constrained framework for recovering directed SC *in vivo*, linking gene expression, anatomical directionality, and large-scale functional hierarchy within one unified model. We found that our genetically informed structure-function model accurately estimates directed structural connections, using only a few gene gradients, across species where ground-truth directed connectivity is available (*C. elegans*, mouse and macaque). In humans, we found corticothalamic feedback connectivity from primary sensory areas, aligning with prior work in non-humans. Human gene ontology analyses suggest a biological mechanism for directionality mediated through molecular gradients. Finally, our model additionally implies a *functional* directionality that recapitulates the traditional unimodal-multimodal functional gradient [56], suggesting that this gradient may emerge from directional signal flow through SC.

## Results

To infer structural directionality, we developed a model that constrains regional source–sink organization to a low-dimensional basis of gene co-expression gradients (Fig. 1). A weighted linear combination of these gradients defines a latent source–sink bias for each brain region we term the “asymmetry metric.” This metric transforms an undirected structural connectome (SC) into a directed SC while preserving the geometric mean between outgoing and incoming edges (Eq. 1). We infer the gradient weights, together with three higher-order network diffusion parameters (Eq. 4), by fitting the model-implied steady-state covariance to the empirical zero-lag functional connectivity matrix through the Lyapunov equation (see Methods, Eq. 6). Thus, the model estimates structural directionality from gene-expression gradients and symmetric structural and functional connectivity, without fitting to an empirical measure of functional directionality.

**Figure 1:**
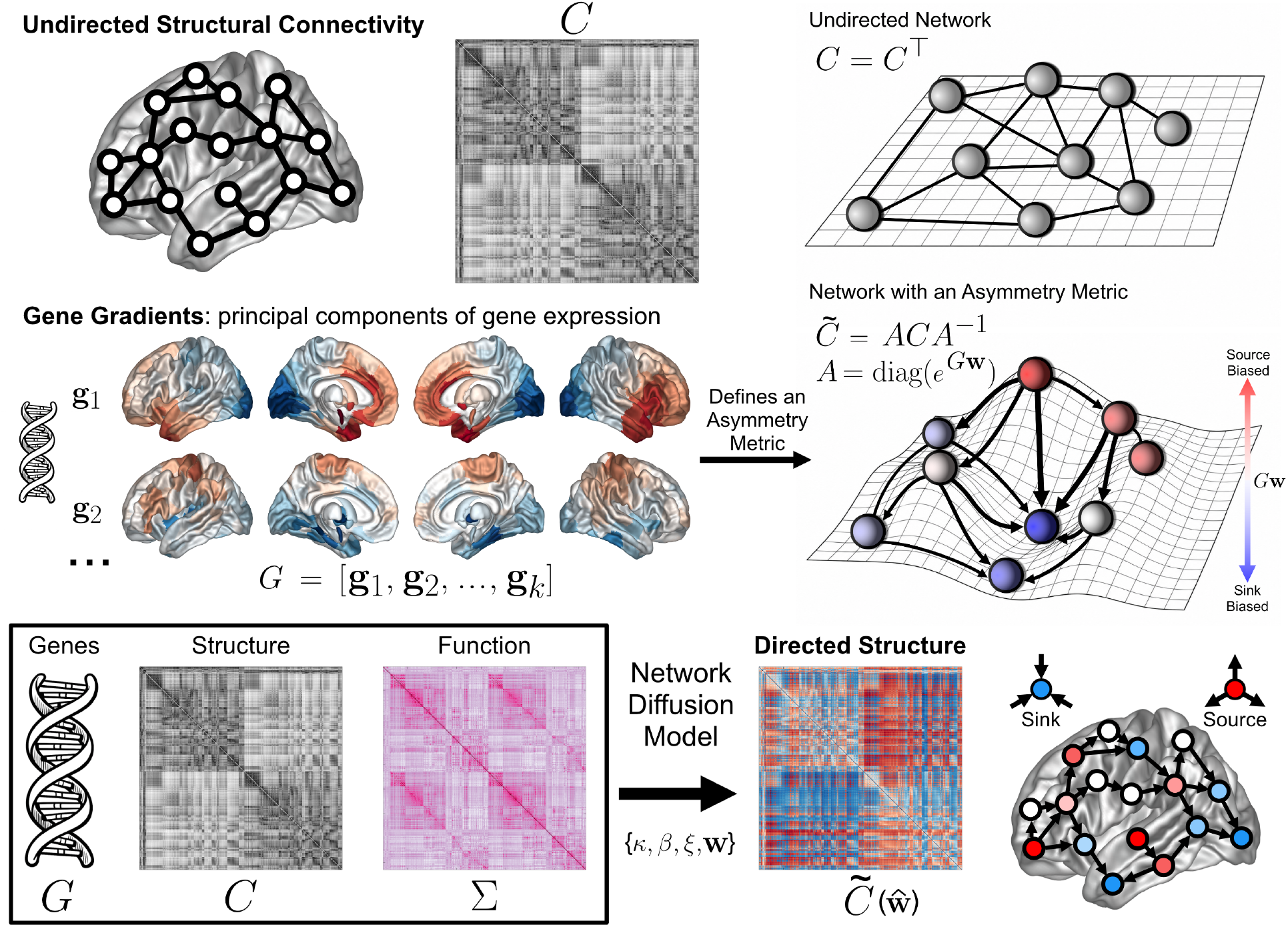
Method Overview. We estimate a node-level *asymmetry metric* using a linear combination of gene gradients (*G***w**). This asymmetry metric is assembled into an asymmetry matrix (*A*), which is used in a similarity transform on the undirected structural connectivity (*C*) to produce our directed SC (*C̃*). This can be intuitively imagined as giving each node an ‘elevation’ on a landscape, resulting in a net directional bias along each edge. Our higher-order network diffusion (HONeD) model has three global parameters {*κ, β, ξ*} and *k* gene weights **w** ∈ R*^k^* that we optimize together to fit to observed functional connectivity (Σ). The final predicted SC uses the optimal gene gradient weights: *C̃*(**ŵ**). Throughout, we follow the convention that network sinks are blue while sources are red.

We first validated our directionality inference approach against ground-truth connectomes from three species: *C. elegans*, mouse, and macaque. We then applied the same approach to infer human structural directionality in the Schaefer-400 atlas with 14 subcortical regions [72, 65], using test–retest fingerprinting to identify the optimal number of gene gradients. We examined the model-predicted human directed SC in terms of regional sinks/sources and directional relationships between resting-state networks. Using the estimated directed SC, we then derived *angular flow* (AF), a model-implied directed functional connectivity constrained by SC. We compared AF with lagged-FC, regression dynamic causal modeling (rDCM), and Granger causality (GC) estimated from human fMRI [28, 10]. Finally, we assessed the relationship between AF net flow and the principal gradient of undirected functional connectivity [56].

### Gene co-expression gradients predict directionality in non-human species

To validate against our three non-human species, we symmetrized the empirical ground-truth directed connectomes, 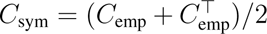, and entered this symmetric matrix, species-specific gene gradients, and the symmetric functional connectivity data into our model. We then compared only the skew (i.e., purely directional) components of the empirical and predicted structural networks, *C*_skew_ = (*C* − *C*^⊤^)*/*2.

Model-predicted directionality significantly correlated with ground-truth directed edges in all three species. Our model-predicted directionality correlated with neuron-to-neuron synaptic directionality in *C. elegans* (*r* = 0.70), tracer-based directionality in mouse (*r* = 0.57), and tracer-based directionality in macaque (*r* = 0.46) (all *p* ≪ 0.001, Fig. 2 a, c, & e). This correspondence remained significant under rank correlation and when evaluating network skew-degrees (Fig. 2 b, d, & f, see Supplemental Section 1 for exact statistics). We found that our model predicted edge-level directionality similarly well within and between hemispheres in *C. elegans* (*r*_ipsi_ = 0.64, *r*_contra_ = 0.57) and in mouse (*r*_ipsi_ = 0.57, *r*_contra_ = 0.59). However, for macaque directionality, only ipsilateral connections were accurate while contralateral connections, which were far fewer in number, were predicted in the opposite direction (*r*_ipsi_ = 0.64, *r*_contra_ = −0.55). We note that the macaque data may have significant limitations, including poor spatial resolution and very few gene expression maps (see Methods), which together may influence the observed results [54]. Nevertheless, these cross-species validations provide strong evidence that the model recovers substantial directional organization under favorable data conditions, while highlighting possible limitations in estimating contralateral directionality from sparse or incomplete data.

**Figure 2:**
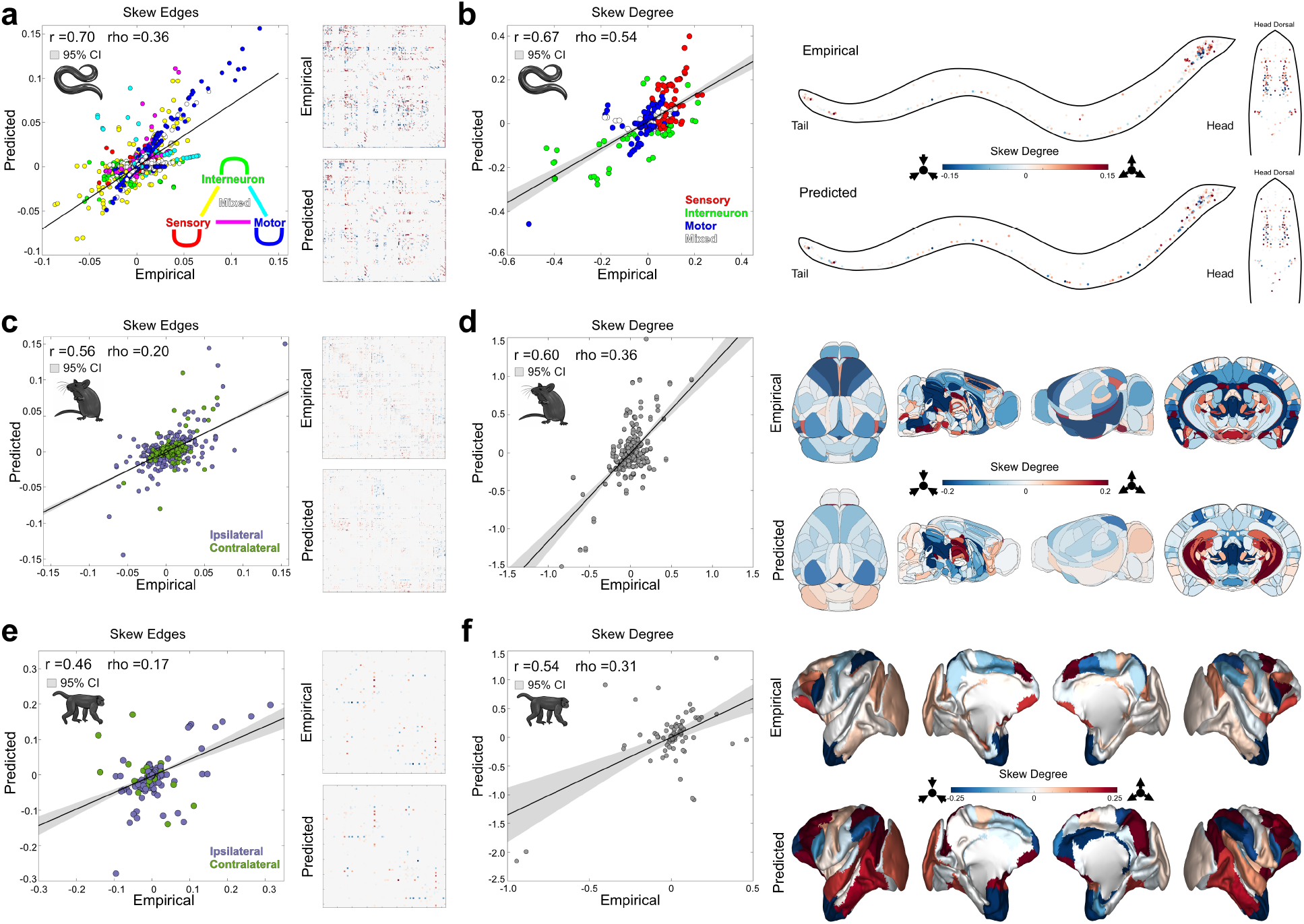
Cross-Species Validation. Comparison of directionality model predictions against ground-truth directed connectivity from three species: *C. elegans* (a & b), Mouse (c & d), and Macaque (e & f). For each species, we show the correlation between observed and predicted skew edges (a, c, e), and the corresponding skew connectivity matrices to the right of the scatter plots. For *C. elegans*, we color each connection by whether the neuron function is sensory (red), motor (blue), interneuron (green), or mixed-identity (white). When connections linked neurons of different types, we colored that connection as an intermediate color between the base colors. For mouse and macaque comparisons, we colored ipsilateral edges purple and contralateral edges dark green. We additionally show the skew degrees (b, d, f), which are the row sums of the skew connectivity matrix (*D*_skew_ = *D*_out_ − *D*_in_), both as a scatter plot and plotted in brain/body spaces on the right. For each scatter plot, we provide both the Pearson r and Spearman *ρ* correlations, all highly significant (*p* ≪ 0.001), as well as the linear-regression best-fit lines with a shaded 95% confidence interval. We show the skew edge correlation results across different numbers of gene gradients in Supplemental Fig. S 6. We also show the empirical SC and FC used to fit the model in Supplemental Fig. S 7.

In our cross-species study the optimal number of gene gradients differed across species (*C. elegans*: *k* = 3; Mouse: *k* = 5; Macaque: *k* = 1; Supplemental Fig. 6), each corresponding to different amounts of explained variance (*C. elegans*: 13%; Mouse: 44%; Macaque: 36%; Supplemental Fig. 6).

### Model-inferred human structural directionality

Because human structural directionality lacks direct ground-truth, we chose the number of gene gradients based on a test-retest fingerprinting analysis of the predicted asymmetry metric (*G***w**). This approach assumes that directed SC estimates should be both replicable within subjects and discriminative between subjects. We evaluated test-retest fingerprinting for 34 HCP test-retest subjects, each with two separate visits, and 724 HCP subjects by splitting resting-state fMRI scans into two ≈ 30-minute blocks. We found that the asymmetry metric had the best test-retest reliability when using *k* = 5 gene gradients (ICC = 0.51 ± 0.03, Discriminability = 0.77, Supplemental Fig. S 8), which explained 59% of gene co-expression variance (Supplemental Section 1). Our model predictions at *k* = 5 gene gradients also replicated in an external dataset (*r* = 0.63*, p <* 0.01, Supplemental Fig. S 9) [69]. Therefore, our subsequent results report the model with *k* = 5 gene gradients.

We show in Fig. 3a the model-inferred group-averaged skew-directionality. We found that many RSNs—Control, Salience, Dorsal Attention, Somatomotor, and Visual—were all strongly directed toward the subcortex (Fig. 3b); conversely, the subcortex had the strongest inter-RSN directionality toward the Limbic network. Across subjects, the Somatomotor and Salience Ventral Attention networks were prominent sources, while the Visual and Control networks showed a more modest source-bias (Fig. 3c). In contrast, the Limbic and Default networks were significantly sink-biased (Fig. 3c). The Subcortical and Dorsal Attention networks showed pronounced directional incoming and outgoing edges, with no significant directional bias at the group level (Fig. 3c).

**Figure 3:**
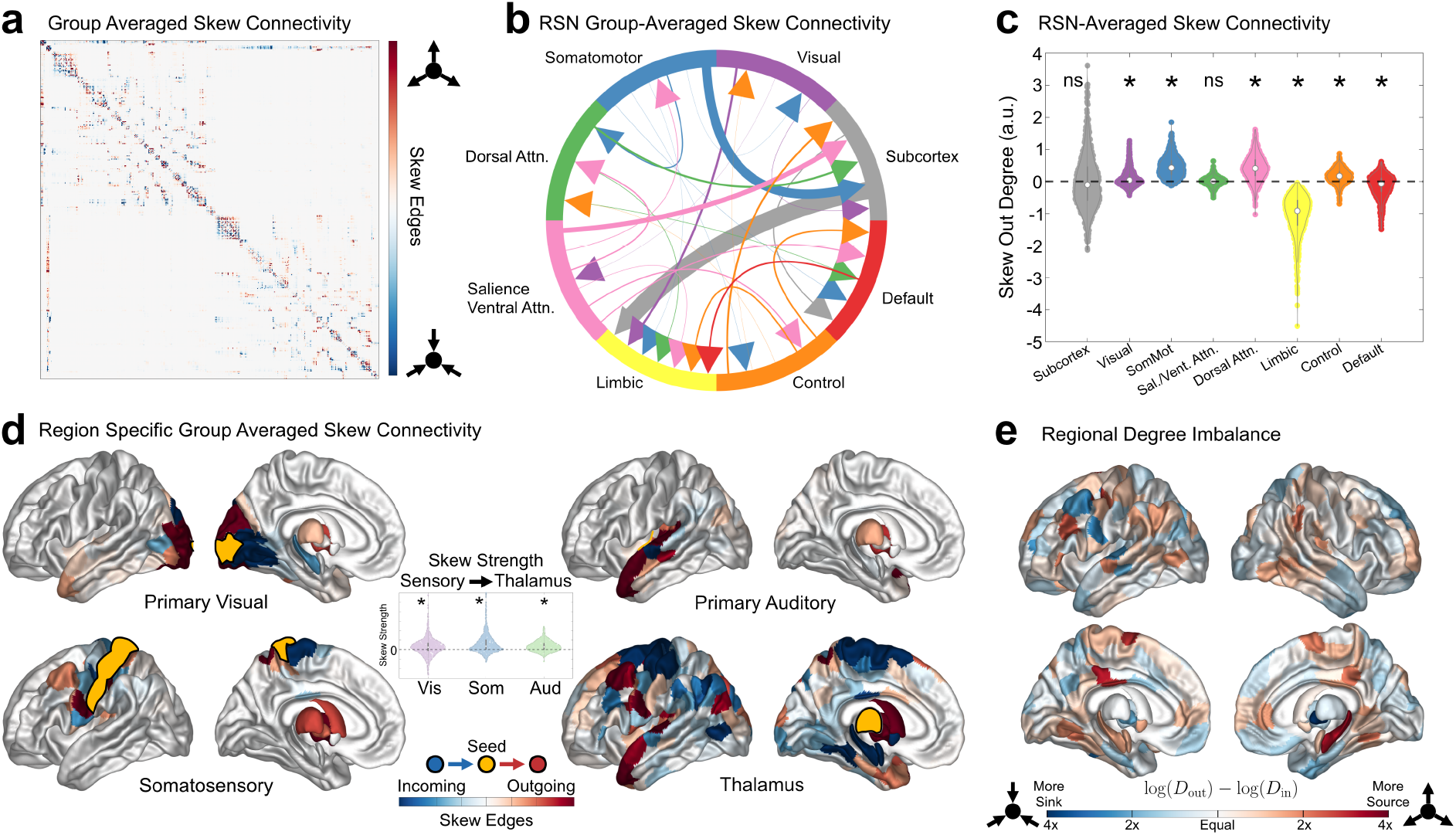
Human Estimated Structural Directionality. (a) The skew component of the group-averaged predicted SC. (b) Significant skew connectivity between resting-state networks. (c) Group-level skew out-degree of each RSN. (d) Select rows from the skew connectome for the primary sensory areas and thalamus showing directional bias between primary and higher-order processing areas. The yellow ROI indicates the exact region(s) we evaluated. The inset shows the group-level weights from each primary sensory area to the thalamus. The primary auditory area is located on Heschl’s Gyrus. All distributions were significantly positive, indicating net anatomical feedback connectivity to the thalamus. (e) The overall degree imbalance for each brain region from the group-averaged SC. The color map is on a log_2_ scale reflecting a multiplicative difference in out-versus in-degrees. Statistical tests were evaluated with a one-way t-test, where *:*p* ≪ 0.01, Bonferroni corrected. We show the fingerprinting and external dataset robustness results in the Supplemental Figs. S 8 and S 9, respectively.

Work in animal models has suggested that anatomical feedback connectivity from primary sensory areas to the thalamus largely outnumbers anatomical feedforward pathways from thalamus to sensory cortices [16, 4]. Therefore, we sought to evaluate and quantify our model-inferred directionality in each of three primary sensory cortices: visual, somatosensory, and auditory. Our model’s predictions align with this previous work, showing group-level bias in anatomical pathways from all three primary sensory areas to the thalamus (Fig. 3d, *p* ≪ 0.001). Additionally, each primary sensory area had distinct patterns of connectivity to other nearby areas. Primary visual areas showed both feedback and feedforward connectivity to areas of both dorsal and ventral visual streams (Fig. 3d). Primary visual areas also showed significant incoming connections from the hippocampus (Fig. 3d). The somatosensory cortex had incoming connectivity from nearby motor areas and outgoing connectivity to higher-level frontal areas (Fig. 3d). The primary auditory cortex on Heschl’s Gyrus showed nearly the reverse pattern: stronger outgoing connectivity to nearby regions, and more distant incoming connectivity from higher-level areas (Fig. 3d). Finally, the thalamus displayed strong directional preferences throughout the cortex and subcortex, which appear suggestive of corticothalamocortical circuits [76].

While the above analyses assess the skew connectivity, measuring each node’s degree imbalance is a way of quantifying the overall multiplicative difference between connections leaving versus entering the region. Across the brain, we found that most regions varied in their degree imbalance within a factor of 2, with some regions, including the hippocampus and pallidum, having as much as a 4-fold degree imbalance (Fig. 3e). The strongest network sources included the hippocampus, medial fusiform gyri, primary motor cortices, and the posterior cingulate; the strongest network sinks included the pallidum and middle frontal gyrus (Fig. 3e, Supplemental Table 1). Although the magnitude of directionality differed, there remained a qualitative symmetry between homologous regions.

### Sinks and sources suggest distinct gene ontologies

To assess the biological plausibility of our model-inferred directed human SC, we investigated whether our model-predicted asymmetry metric implicated genes with interpretable ontologies. We calculated the cosine similarity between our consensus-fit asymmetry metric (*G***w***_c_*) and each of the 15, 633 gene expression maps. Using the top 1, 000 genes aligned with sinks or sources, respectively, we evaluated their gene ontologies with statistical comparisons to both our pool of 15, 633 genes and a null distribution defined by a spatial-autocorrelation (SA)-preserving Moran spectral randomization of the empirical map [17, 48] (see Methods for details). We calculated the “Enrichment Ratio” as the empirical number of genes related to the gene ontology term normalized by the mean number of genes across all SA-null maps associated with the same term.

We found that sink- and source-aligned genes were each associated with different gene ontology terms. Sink-aligned genes were enriched for ontologies related to protein synthesis, translation, and endoplasmic reticulum (ER) processing and quality control (Fig. 4c). The sink-axis was also enriched with genes related broadly to neural development and axon guidance, especially the regulation of SLIT and ROBO expression, as well as immune and broad disease- and infection-associated pathways (Fig. 4c). In contrast, source-aligned genes showed significant enrichment for ontology profiles related to cellular structural constituents, including filaments and keratinization, as well as chemosensory receptor activity and G protein-coupled receptor signaling (Fig. 4c). Together, these results suggest a model-informed hypothesis in which sink-aligned regions are enriched for molecular programs related to protein processing and developmental guidance cues, whereas source-aligned regions are enriched for cytoskeletal and receptor-related programs that may support cellular responsiveness to such molecular landscapes.

**Figure 4:**
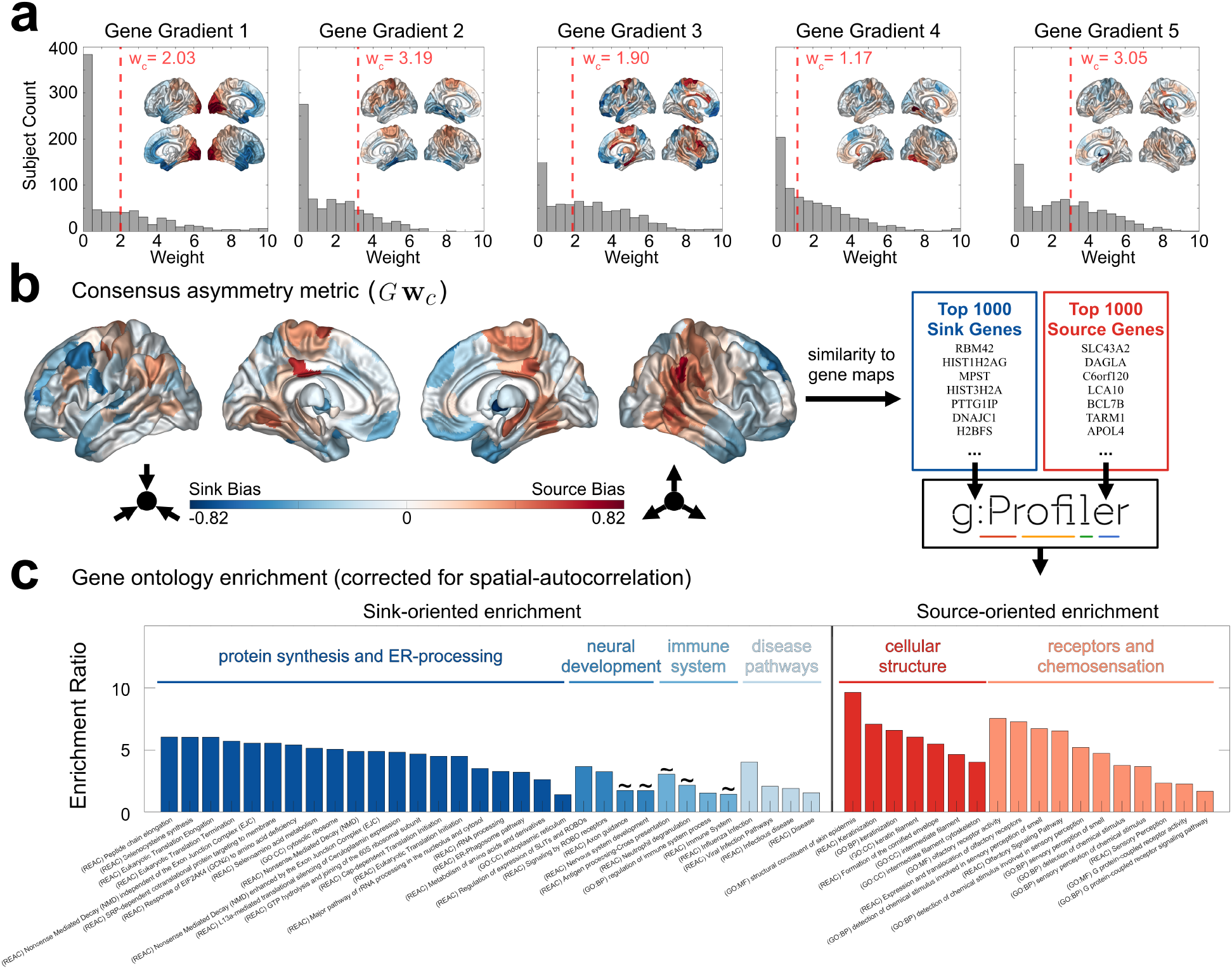
Model-Predicted Gene Ontologies. (a) The model-predicted gene gradient weights across all subjects (gray) and the consensus-fit gene gradient weights (dashed red line). Insets show each respective gene gradient. (b) The model-predicted asymmetry metric for the consensus fit (i.e., the linear combination of gene gradients, *G***w***_c_*). Blue regions are sink biased, red regions are source biased. We identified the 1000 most sink- and source-aligned genes for input to g:Profiler. (c) A gene ontology enrichment analysis, corrected for spatial autocorrelation. We show all FDR-corrected significant gene ontology terms for the sinks (blue) or sources (red). Subsequent statistical testing compared these terms against spatial-autocorrelation (SA) preserving null models. The “Enrichment Ratio” is the empirical number of genes related to the term normalized by the mean number of genes across all SA-null maps associated with the same term (Equation 12). Gene ontology terms with a ∼ were statistically significant (FDR *p <* 0.05) under the g:Profiler general ontology analysis, but trending (0.05 *< p*_SA_ *<* 0.1) under the SA-null distribution; all other terms were significant under both.

### Angular flow relates directed structure to directed functio

As we show in Methods, we can use our directed SC operator to calculate a measure of directed flow through the brain, which is related to the brain’s state-space angular momentum (Fig. 5a). Accordingly, we term this measure “angular flow” (AF). We assessed whether AF offers complementary directional information to conventional causal discovery metrics in humans by comparing with lagged-FC, regression dynamic causal modeling (rDCM), and Granger causality (GC). To keep these metrics well-posed and tractable, we ran our comparisons in the Schaefer-100 atlas with 14 subcortical regions.

**Figure 5:**
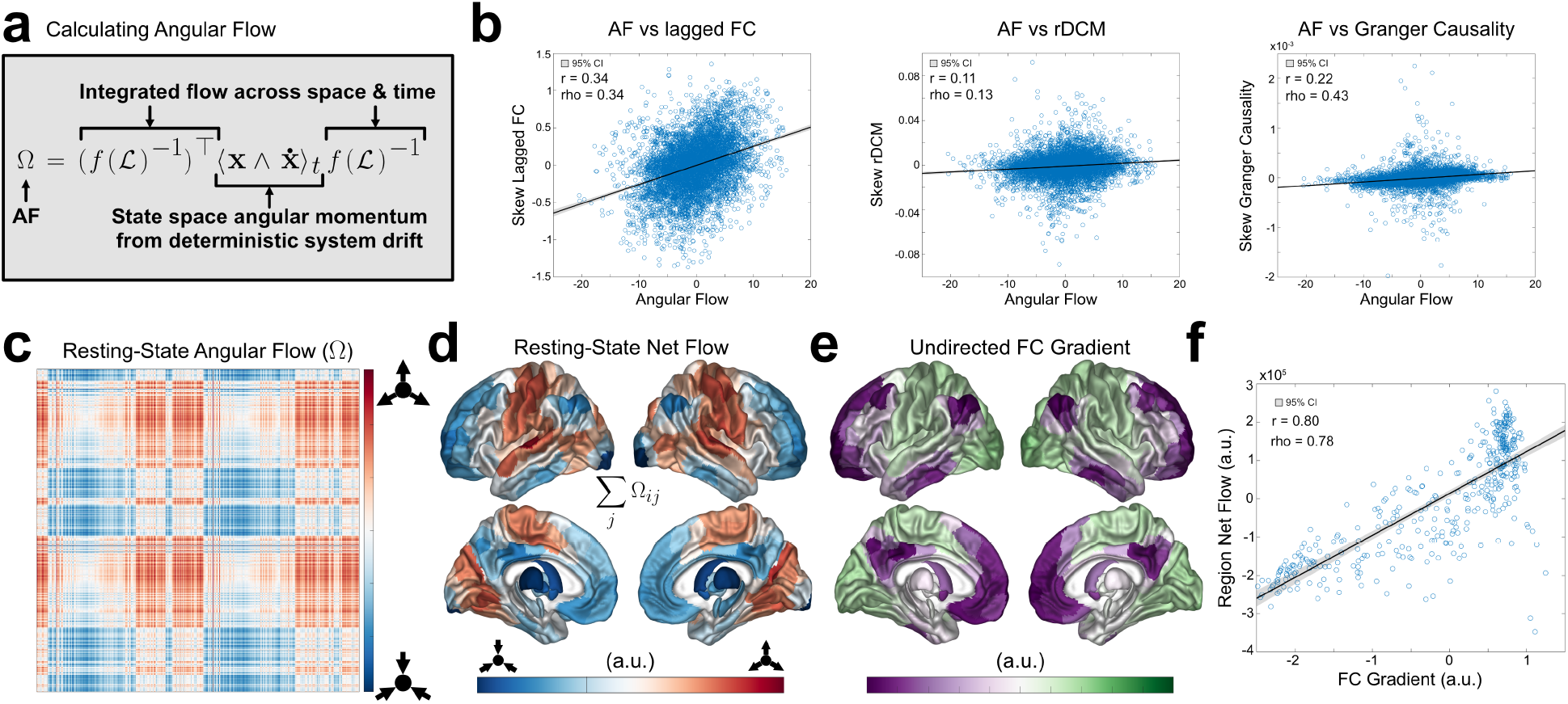
Angular Flow. (a) The AF equation, describing signal flow governed by the HONeD model *f* (L) and its relationship to probability angular momentum ⟨**x**∧**ẋ** ⟩*_t_*. (b) The comparisons between AF and (left-to-right) the human lagged-FC, rDCM, and GC, respectively (all in the Schaefer-100 atlas). (c) The angular flow (AF) during resting-state fMRI, averaged across subjects. (d) The sum along each row of AF to obtain the network’s net flow. (e) The first principal gradient of the group-averaged functional connectivity. (f) A scatter plot comparing (d) with (e), showing a highly significant correlation (*r* = 0.8*, p*_SA_ ≪ 0.001). The spatial autocorrelation preserving null-distribution is shown in Supplemental Fig. 11; the underlying diffusion operator is shown in Supplemental Fig. 12.

We found that all three causal discovery methods significantly correlated with AF (Lagged-FC: *r* = 0.34; rDCM: *r* = 0.11; GC: *r* = 0.22; all *p* ≪ 0.001; Fig. 5b), and these results remained significant under rank correlation (see Supplemental Section 1). This provides convergent evidence that AF captures a broad directional bias shared, albeit weakly, by other causal discovery methods, but not reducible to any of them.

### Angular flow recapitulates the principal FC gradient

AF during resting-state fMRI showed distinct sink/source relationships between RSNs. Source RSNs consisted of the visual, somatomotor, dorsal attention, and salience/ventral attention networks; meanwhile, sink RSNs consisted of the default, control, limbic, and subcortical networks (Fig. 5c-e). Moreover, AF had regional net flow that strongly correlated with the principal functional connectivity gradient (*r* = 0.80*, p*_SA_ ≪ 0.001; Fig. 5f) [56]. Since the inverse of our HONeD operator generates the flow dynamics, we investigated whether this correspondence to the FC gradient was trivially related to the inverse operator; however that was not the case (*r* = 0.04, Supplemental Fig. 12). This emphasizes that the emergence of the principal gradient results from a combination of the inverse diffusion operator and the empirical signal covariance. In principle, AF could also be calculated on an undirected SC, which we found also has a correspondence with the FC gradient, albeit significantly weaker (*p <* 0.001; Supplemental Fig. 13). Additionally, we compared the FC gradient to empirical lagged-FC, finding that lagged-FC showed substantially weaker alignment with the FC gradient than AF (*p* ≪ 0.001; Supplemental Fig. 13). Together, these results highlight how AF calculated with the directed SC recapitulates the principal FC gradient.

## Discussion

Inferring directed structural connectivity (SC) at the macroscale remains a central challenge in network neuroscience. Here, we constrain directionality to a low-dimensional basis of gene coexpression gradients and fit their contributions using a structure-function model. The model recovered significant directional organization across three non-human species (Fig. 2), and, in humans, produced reliable subject-specific estimates with biologically interpretable sink/source organization and gene ontologies (Figs. 3 & 4). The model operator also implied a measure of functional “angular flow” (AF) through SC that recapitulates the principal gradient of functional connectivity (Fig. 5). Together, these findings link regional gene expression, anatomical directionality, and large-scale signal flow within a unified framework.

### Gene gradients constrain structural directionality

A central implication of these results is that gene co-expression gradients define a low-dimensional biological manifold for network directionality. Although prior work has suggested links between genes and structural connectivity, it has generally lacked a model to relate them [26, 50, 6, 20, 85]. Here, we provide such a genetically informed structure-function model, which suggests that genes impose a directional bias at both neuronal and regional levels. Past work on structure-function modeling has argued that lagged functional connectivity is necessary for SC directionality to be identifiable [31]. However, our results point to gene expression as a parsimonious and biologically plausible constraint from which directional properties can emerge even for zero-lag covariance. This constraint collapses an otherwise underdetermined *n*^2^-dimensional edge-direction problem into a low-dimensional node-level parameterization, making directionality estimation tractable and empirically testable.

### The structure-function relationship reveals latent directionality across scales

The present work adds to a growing literature showing that structural information can impose strong constraints on functional organization, especially within linear models [1, 2, 14, 82, 62]. That our model can approximate ground-truth directed SC by fitting to undirected functional connectivity supports the broader view that a *directed* SC provides a macroscopically linear scaffold for brain function with observable consequences.

Somewhat surprisingly, we found that our linear graph diffusion model was effective in fitting to *C. elegans* data and accurately recovering the underlying synaptic directionality (Fig. 2). This suggests that even a relatively small network of neurons with highly nonlinear dynamics may still have a form of macroscopic linearity that our model can observe. This is especially notable because previous work studying signal propagation in *C. elegans* showed that SC poorly predicted functional signaling when using a neural mass model [67]. Although the present work does not yet fully resolve the SC-FC misalignment tension in *C. elegans*, these findings suggest the potential for linear structure-function relationships in the brain to remain relevant even within relatively small neural populations.

### Model-predicted gene ontologies have biological interpretability

Our model predicted sink-aligned regions that were enriched for genes involved in protein synthesis and neural development while source-aligned regions were enriched for cellular structure and chemosensory receptor programs (Fig. 4). One especially interesting result was the model-predicted enrichment for SLIT and ROBO expression. SLIT ligands and their ROBO receptors are well documented guidance systems that regulate axon attraction and repulsion in the developing brain [87, 53, 79, 71, 57]. Meanwhile, intermediate filaments are broadly involved in cell and tissue mechanics, contributing to mechanical stability, cytoarchitecture, cell-shape regulation/remodeling, and cell migration [42, 51], providing a biological interpretation for their enrichment in source-aligned regions. Source-aligned regions were also enriched for chemosensory olfactory receptors (ORs), which are known to guide axon growth in olfactory circuits [84, 21]. ORs also show significant ectopic expression during development [47], which, when combined with our results, suggests ORs may have a more general role in neural development than currently understood. Contextualized by this literature, our model hypothesizes that sink-aligned regions may contribute molecular guidance landscapes, while source-aligned regions may be enriched in receptor and cell structure programs that support sensing and responding to such landscapes. However, given a lack of ground-truth in humans, it is important to emphasize that these interpretations should be viewed cautiously as model-driven hypotheses to inspire future experiments rather than decisive results.

### Model-predicted directionality aligns with known anatomical pathways

Our human directed SC prediction appears consistent with well-studied anatomical pathways in non-humans. There is considerable scientific discussion on the role and quantity of feedback versus feedforward connectivity between sensory areas and the thalamus [16, 4]. While corticothalamic connectivity is generally reciprocal, evidence suggests a bias toward more corticothalamic projections across sensory modalities [61, 95, 102, 68]. Our model similarly showed corticothalamic bias in primary sensory areas, and is therefore broadly consistent with prior non-human anatomy literature (Fig. 3). Feedback connectivity from higher-level sensory areas to primary sensory areas is also well documented [59, 49], and was also observed in our model predictions (Fig. 3). However, our model did not simply predict the thalamus as a global sink, but instead predicted patterned bidirectional connections throughout the cortex, suggestive of corticothalamocortical circuits [76]. At the network level, the limbic RSN was the most decisive sink across subjects, which aligns with previous hypotheses that these regions are targets of top-down allostasis-driven predictions [11]. While it is reassuring that the model appears generally aligned with previous research, it is crucial not to over-interpret model predictions and instead recognize that these predictions are merely hypotheses to guide future empirical work.

### Angular flow as a new measure of functional directionality

Most estimates of directionality in human neuroimaging infer effective connectivity from interregional predictability, as in Granger causality [74, 10], or from model-based approaches such as dynamic causal modeling [25]. These approaches become computationally difficult and increasingly ill-posed at whole-brain scales [25], and they generally do not incorporate information from the weighted structural connectome.

Angular flow (AF) provides a complementary, structure-informed description of directional dynamics. Rather than estimating pairwise causal influence, AF analytically integrates the fitted HONeD dynamics with the full weighted SC to quantify model-implied signal flow between regions. AF was modestly correlated with lagged-FC, rDCM, and multivariate Granger causality, indicating that these measures capture partially overlapping directional information despite their distinct interpretations (Fig. 5). Importantly, AF distinguishes the directionality of the anatomical scaffold from the dynamics unfolding upon it. For example, the thalamus is only weakly sink-aligned in the modeled SC, but it becomes the strongest sink in AF. Thus, dynamical flow between brain regions need not be fully reduced to the pairwise anatomical directionality, but emerges from the interaction between network architecture, system dynamics, and the presence of exogenous or endogenous drive.

The first-principles derivation of AF also reveals a surprising relationship to the brain signal’s state-space angular momentum. In non-equilibrium statistical mechanics, the time-averaged quantity ⟨**x** ∧ **ẋ** ⟩*_t_* is termed “probability angular momentum” and characterizes persistent state-space circulation, time-reversal asymmetry, and broken detailed balance [27, 106, 107, 100, 36, 78]. AF is distinct from probability angular momentum because the HONeD operator defines the dynamical estimate of **ẋ**, and the inverse operator transforms the resulting circulation into a pairwise flow between regions. Although probability angular momentum has been used to characterize non-equilibrium climate oscillations [106, 100], to our knowledge, it has not previously been explicitly applied to empirical brain activity.

### The principal FC gradient emerges as the angular flow through directed SC

AF may explain the emergence of the unimodal-multimodal principal functional gradient in undirected functional connectivity. Since its first description by Margulies et al. [56], much work has emphasized the primacy of this unimodal-multimodal gradient in the brain [41]. Previous work has related this principal gradient to the brain’s structure-function relationship, even when SC-FC coupling is defined in several distinct ways [64, 66, 13, 98]. AF extends and refines these insights by suggesting that the unimodal-multimodal gradient arises from the net flow of brain signals through the directed SC from unimodal to multimodal cortical areas (Fig. 5). Importantly, lagged-FC failed to recover this relationship (Supplemental Fig. 13), emphasizing the role of the structure-function model in revealing this correspondence. This result dovetails with our prior work on network deconvolution [80]: when passive network diffusion over SC is removed from fMRI, functional gradients become remodeled; when directional diffusion is estimated on empirical brain signals, the functional gradient reappears as the pattern of net flow through structure.

### Limitations and future work

These results should be interpreted with appropriate caution. Human directed SC here remains a model prediction rather than a directly observed anatomical measure. The non-human validations are limited by the availability of only one consensus ground-truth per species, and those groundtruth datasets differ in completeness, scale, and biological meaning. The model also depends on how regional gene expression maps are estimated, including which parcellation is chosen, which genes are available, and how missing tissue is interpolated. These ground-truth limitations were especially relevant for the macaque data, where directionality was consolidated by CoCoMac across many separate studies and both the gene spatial coverage and diversity were highly limited. We speculate that these issues may explain why the macaque model only used one genetic gradient, and why the prediction itself was the weakest of the three ground-truth analyses.

Methodologically, our directionality parameterization is effective but still bespoke: a diagonal similarity transform is mathematically convenient and biologically suggestive, yet it is not itself a mechanistic developmental model of axonal targeting. Likewise, the fitted diffusion operator is linear and stable, which likely restricts the space of recoverable directed dynamics even as it guards against overfitting. Our AF measure inherits these assumptions and introduces an additional interpretational caveat: an inhibitory influence between two regions may be partially conflated with a reversed directionality. Future work should therefore consider testing alternative generative mechanisms for genetically influenced asymmetry and assessing the behavioral relevance of task-based changes in AF.

## Conclusion

We introduced a genetically informed structure-function model for recovering directed structural connectivity and validated it against ground-truth connectomes in *C. elegans*, mouse, and macaque. Applied to humans, the model revealed biologically interpretable source–sink organization and enabled a structurally informed measure of directed functional flow, termed angular flow (AF). AF recapitulated the principal functional connectivity gradient, linking molecular organization, anatomical directionality, and large-scale functional hierarchy within a unified framework. Overall, this framework provides a fundamentally new way to estimate and analyze both directed structural and functional connectivity in the brain.

## Methods

### Theory

Here, we describe the structure-function model used to predict directed structural connectivity. In essence, our model predicts that resting-state functional covariance is close to a stationary covariance, which we formalize using the Lyapunov Equation within a multivariate Ornstein–Uhlenbeck (MOU) process. The model operator that governs the brain’s internal dynamics over the structural connectome is a higher-order (i.e., polysynaptic) network diffusion model. Our goal is to find a node-level directionality bias, parameterized by gene gradients, that optimizes this stationary structure-function alignment. In the following, we construct this structure-function model by first parameterizing structural directionality with gene gradients, then introducing the higher-order graph Laplacian operator, and finally relating the resulting dynamics to stationary empirical covariance.

### Parameterizing Directionality with Gene Gradients

We first obtained gene expression maps (*E* ∈ R*^m^*^×*n*^) for all *n* brain regions (or neurons in the *C. elegans* case), z-scored each map, then calculated the principal components as the eigen-decomposition of the gene expression covariance matrix (*E*^⊤^*E* = *G*Λ*_E_G*^⊤^). Hereafter, we term these principal components “gene gradients”. Using the gene gradients associated with the *k* largest eigenvalues (*G*^(*k*)^ ∈ R*^n^*^×*k*^), we imposed directionality by constructing a node-wise *asymmetry metric* as a linear combination of genetic gradient basis vectors with weights (**w** ∈ R*^k^*^×1^): *G*^(*k*)^**w**. This asymmetry metric is built into a diagonal positive-definite matrix 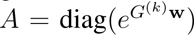 which induces a similarity transform on our undirected structural connectivity matrix *C* ∈ R*^n^*^×^*^n^*:

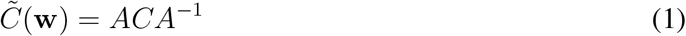

Element-wise, this is 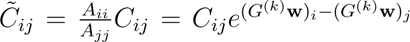, making it clear that each entry in *C* is multiplied by an asymmetry factor given by the diagonal of *A*^1^. This transformation preserves the geometric mean of reciprocal edge weights, and it also preserves all non-zero edges (i.e., the model will not add new edges that were not present in the original connectome). Throughout, we follow the convention that *C̃_ij_* indicates a directed connection from node *i* to node *j*. In the directed network *C̃*, we may calculate the *out-degree* (*D*_out_ = diag(Σ*_j_* C̃*_ij_*)) and the *in-degree* (*D*_in_ = diag(Σ*_i_* C̃*_ij_*)) representing how strongly that region transmits or receives connections, respectively.

#### Directed Graph Laplacian

The graph Laplacian is a graph analogue to the Laplace-Beltrami operator for manifolds [46]. It is critical to modeling how signals move through networks, and it is the heart of graph diffusion models. Here, we chose to use the column-stochastic Laplacian:

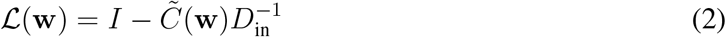

The graph Laplacian normalizes each node by the network’s in-degree, thereby assuming that each brain region changes its signal based on the weighted average of all incoming signals. We chose this specific Laplacian normalization based on previous work demonstrating that in-degree normalized networks outperform other normalization strategies when predicting directed functional connectivity [82].

#### The Higher-Order Network Diffusion (HONeD) Model

Prior work, including our own, has generalized the network diffusion model on networks to account for multi-step connections through SC [80, 14, 82]. Here, we again leverage higher-order network diffusion (HONeD) as our structure-function model operator to identify SC directionality. We define our diffusion operator as:

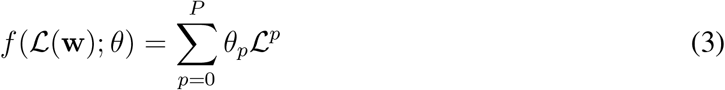

Here, *P* is the order of the Taylor series approximation, and L is the directed graph Laplacian of structural connectivity. As in past work, we chose to limit the network diffusion model to two steps (L^2^) through the network^2^ [80, 14]. Therefore, the HONeD operator for the present work becomes

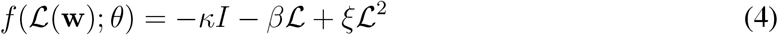

Here, we denote the terms from the Taylor expansion *θ*_0_ = −*κ*, *θ*_1_ = −*β*, and *θ*_2_ = *ξ* for ease of referencing. Written in this way, *κ*, *β*, and *ξ* are all constrained to maintain model stability and identifiability. In the following, we use *f* (L) as a shorthand for this parameterized function of the graph Laplacian.

#### Predicting Structural Directionality with HONeD

Network diffusion may be well-suited to identifying directionality since its influence on brain signal covariance should remain stable across long time scales. HONeD can be expressed as the drift operator of a structurally constrained multivariate Ornstein–Uhlenbeck (MOU) process, or equivalently a linear time-invariant (LTI) graph-diffusion system driven by stochastic input [31, 33, 1, 93]. In this model, we consider a brain signal **x**(*t*) ∈ R*^n^* with an actuation matrix *B* ∈ R*^n^*^×*ℓ*^ and a stochastic driving signal 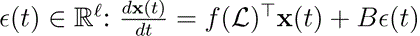3.

This MOU process determines the evolution of the signal covariance Σ(*t*) = E[**x**(*t*)**x**(*t*)^⊤^] through the Lyapunov equation [8]:

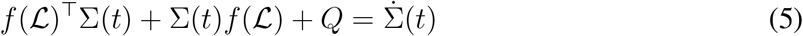

Commonly, the Lyapunov equation is solved at steady state where Σ̇ (*t*) = 0 denotes the steady-state condition [31]. At steady state, this derivative is zero. We approximate the stationary covariance with the sample covariance 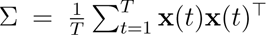, and with the noise covariance *Q* = *B* E[*ɛ*(*t*)*ɛ*(*t*)^⊤^]*B*^⊤^. For a thorough derivation, see Supplemental Section 2. By estimating the sample covariance of brain signals as Σ, we can leverage the Lyapunov equation to define a cost function for our model:

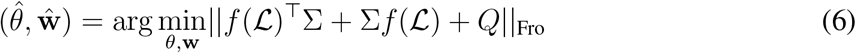

With this cost function, we acknowledge that the brain is not fully stationary, so we instead find the optimal model parameters ({*θ̂*, **ŵ** }) that bring the system as close to a stationary state as possible. When fitting the model, we chose to approximate *Q* using the diagonal of the empirical covariance: diag(Σ). Using these optimal values, we constructed a directed *f* (L) that best describes the observed stationary diffusion process in the brain.

#### Predicting Functional Directionality with Structural Directionality

Once our directed HONeD operator has been fit to empirical data, we can then leverage that same operator with the same parameters to better understand the directionality of functional brain signals.

Following previous work, we approach functional directionality by considering time lags in the system. More precisely, our model predicts that 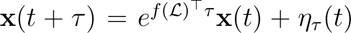, where 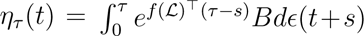 is the future white noise accumulated after time *t*. This yields a prediction for the time-lagged covariance:

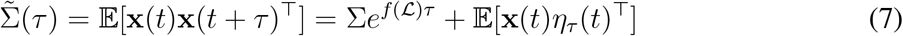

Then Σ̃>(*τ*)*_ij_* quantifies how fluctuations in region *i* at time *t* covary with region *j* at time *t* + *τ*. ^4^ Here, we assume that the white noise is independent of the current state, causing the second term to vanish: E[**x**(*t*)*η_τ_* (*t*)^⊤^] = 0. Now we will define the total flow (*F*) of functional signals through all paths in the structural connectome as the integral across all lags:

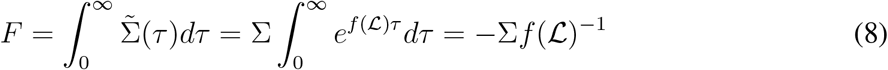

This uses the identity 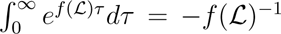, which holds whenever *f* (L) is Hurwitz.^5^ The matrix −*f* (L)^−1^ is the time-integrated evolution operator for the system, and it bears some similarity to previously defined measures of flow based on a different formulation of a linear dynamical MOU process [32]. Therefore, *F* can also be viewed as a form of “flow” in the sense that it casts the empirical covariance (conventionally known as “functional connectivity”) through the directed SC operator integrated *across all time*, reflecting the aggregate flow of signal from any region *i* to region *j* across all possible network paths.

#### Deriving the Brain Signal’s Angular Momentum

In this section, we show that this hybridization between the empirical brain function and the model operator has a fundamental relationship to a form of angular momentum in the brain. To show this relationship, first consider the purely directional, anti-symmetric component of *F* :

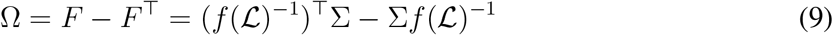

When we factor out the time-integrated operator from both sides, it reveals a relationship to an *angular momentum*-like tensor, or more precisely bivector [44], of the brain signal’s state space:

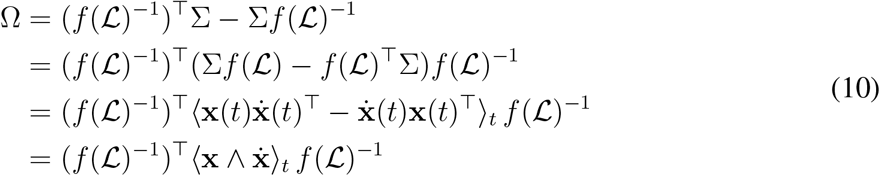

We take **ẋ** (*t*) = *f* (L)^⊤^**x**(*t*) to be the fully deterministic drift without noise. Here, **x** ∧ **ẋ** can be interpreted as an angular momentum bivector, analogous to classical angular momentum in physics [44].^6^ In non-equilibrium statistical mechanics, this time-averaged quantity ⟨**x** ∧ **ẋ** ⟩*_t_* is termed “probability angular momentum” and characterizes persistent state-space circulation, time-reversal asymmetry, and broken detailed balance [27, 106, 107, 100, 36, 78]. Given its origin as the antisymmetric component of covariance-weighted integrated flow, and its formal relationship to a state-space angular momentum bivector, we refer to Ω as *angular flow* (AF). Therefore, AF has the interpretation of transforming the brain signal’s state-space angular momentum through the time-integrated system dynamics.

Using AF, we may calculate the net flow for each brain region as Σ*_j_* Ω*_ij_* = Σ*_j_* (*F_ij_*− *F_ji_*), which is positive if there is net flow moving *out* of a region, and negative if there is net flow *into* a region. We can interpret Ω as an antisymmetric edge-flow field, where Ω*_ij_ >* 0 implies a net flow on that edge pointing *i* → *j*. Since Ω*_ji_*= −Ω*_ij_*, AF can formally be called a graph 1-cochain [3]. In this light, net AF has a natural interpretation of taking the discrete (graph) divergence of this flow field: (div Ω)*_i_*= Σ*_j_* Ω*_ij_*, which is the signed sum of all flows incident on region *i* and therefore measures the total outward minus inward flow.

It is worth emphasizing again that AF is *not* purely an SC-based prediction for functional connectivity, nor is it a trivial reproduction of the original signal covariance (Σ). AF also does *not* imply a notion of pairwise causality; rather, it implies a whole-brain ‘tilt’ of signal flow through the entire network between any two brain regions across time. AF is a quantitative measure that hybridizes the prediction of a directed structural operator governing brain signal flow and the empirically observed signal covariance itself. Since it is derived from the HONeD operator, it inherits the constraints imposed by passive diffusion through SC when calculating the asymmetric flow between regions.

### Non-Human Datasets and Processing

#### Caenorhabditis elegans Dataset

We leveraged the publicly available *C. elegans* dataset provided by `wormatlas`, which includes recordings from 300 neurons imaged from 113 individuals [67]. In particular, it includes a directed neuron-to-neuron structural connectome [101, 103], single-cell transcriptomics for 13, 669 genes [86], and neuron-level Ca^2+^ fluorescence imaging time series where optogenetic stimulation was applied to single cells [67]. The final *C. elegans* connectomes reflected structural synaptic connectivity between 296 cells, where 4 cells were excluded since they lacked any empirically observed synaptic connections. We took the empirical ground-truth neuron-to-neuron structural connectome and symmetrized it as *C*_sym_ = (*C* + *C*^⊤^)*/*2, thereby destroying all directional information, before entering it into our directionality model. For an approximation of the signal covariance between neurons, we used the empirical preprocessed Ca^2+^ imaging time series across all 113 individuals and calculated Σ*_ij_* = ⟨**x***_i_*(*t*)**x***_j_*(*t*)⟩*_t_* ∀*t* | **x***_i_*(*t*), **x***_j_*(*t*) ̸= 0. Since each cell had its own transcriptomic profile, we calculated the principal components of the full matrix and used the *k* leading principal components in our model (see Theory section for mathematical details).

#### Mouse Dataset

The Allen Mouse Brain Connectivity Atlas, the first whole-brain mouse connectome, was published and made publicly available in a landmark study by the Allen Brain Institute [63]. They used an anterograde tracer with 469 image sets to identify directed structural connections between 213 regions across the entire mouse brain. Each region was assessed for either ipsilateral or contralateral connectivity, meaning that left and right homologs were used together in directional estimations. Therefore, to construct a whole-brain connectome with 426 regions to serve as ground-truth, we assumed that left-right homologous regions had equivalent ipsilateral/contralateral directionality. Region-level gene expression data came from the coronal series of the in situ hybridization (ISH)-based Allen Gene Expression Atlas (AGEA) with 4083 genes [52].

Preprocessed functional MRI data were obtained from nine mice (age: 9 ± 0.05 months) with a total of 40 minutes of resting-state fMRI scan time for each mouse [88]. Preprocessed fMRI and anatomical images were provided registered to the Australian Mouse Brain Mapping Consortium (AMBMC) MRI template. We then registered the Allen Brain structural template at 10 *µ*m to the AMBMC mean fMRI volumes using Advanced Normalization Tools (ANTs) with linear interpolation in Python. We then used this transform between spaces to map the Allen Brain atlas with 426 regions to the AMBMC space with generic label interpolation. With the atlas in AMBMC space, we used nilearn to calculate the mean time series for all regions.

#### Macaque Dataset

For the non-human primate dataset, we used an OpenNeuro macaque dataset that includes fMRI time series from nine macaques (eight *Macaca mulatta*, one *Macaca fascicularis*) and a dMRI-derived consensus SC parcellated in the RM atlas with 82 cortical brain regions [75]. Notably, the SC was directed, with directionality coming from aggregating results from tracer-based studies in the CoCoMac dataset [83]. This tracer-based directionality was treated as a ground-truth to which we compared our model performance. When entering this connectome into our directionality model, we symmetrized the macaque connectome (*C_M_*) as 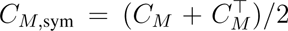. The preprocessed fMRI time series were obtained in a standardized F99 volumetric space after confound regression, and we parcellated the time series by averaging all voxels in each region using the macaque volumetric RM atlas. Macaques were lightly anesthetized before their scanning session, and anesthesia was maintained using 1–1.5% isoflurane during image acquisition [75]. For full acquisition and processing details, see Shen et al. [75]. Region-wise transcriptomics was obtained from a publicly available dataset with 131 genes [54].

#### Statistical Comparisons to Ground-Truth Connectomes

When comparing model predictions to the ground-truth in the above non-human species, we calculated the Pearson correlation between the empirical and predicted skew connectivity matrices, *C*_skew_ = (*C* − *C*^⊤^)*/*2, as well as empirical and predicted skew degrees, *D*_skew_ = *D*_out_ − *D*_in_. We used only non-zero edges since the absence of an empirically reported directed edge does not necessarily reflect a true absence of directed connectivity between observed brain regions. When p-values were below computer precision, we denoted them as *p* ≪ 0.001.

### Human Dataset and Processing

We used 770 subjects from the Human Connectome Project (HCP) Young Adult dataset in this study [94] ^7^. We excluded all HCP subjects that had documented quality control issues. Unprocessed HCP data were downloaded and organized into BIDS format [37]. Anatomical scans were collected with an MPRAGE T1-weighted sequence. Functional MRI (fMRI) data included four 15-minute rest sessions and two task sessions for six tasks (Motor, Language, Gambling, Social, Relational, and Working Memory), all with a repetition time (TR) of 720 ms. Additionally, we used the preprocessed diffusion MRI (dMRI) data made available by HCP, which includes merging high-fidelity directions across multiple multi-shell dMRI acquisitions (b = 1000, 2000, & 3000) and correcting for phase-encoding polarity distortion [34]. We applied a state-of-the-art image processing pipeline, micapipe [19], which processes the anatomical, functional, and diffusion MRI data in a coherent framework to produce subject-specific structural connectivity and functional regional time series in the Schaefer atlas with 400 cortical regions [72], each with connectivity to an additional 14 subcortical regions [65]. Regional expression maps for 15, 633 genes were obtained using the `abagen` toolbox in our same parcellation space with linear distance-based interpolation for any missing regions [58, 40].

The structural connectivity was computed using MRtrix3 and probabilistic tractography, generating 10 million streamlines throughout the gray-white matter interface (maximum tract length = 400, minimum length = 10, cutoff = 0.06, step = 0.5, angle curvature = 22.5^◦^) using the iFOD2 algorithm and 3-tissue anatomically constrained tractography [91, 92, 45, 19]. Tractograms were filtered using SIFT2 [81], reweighting streamlines by cross-sectional multipliers to provide connection densities that are biologically valid measures of fiber connectivity.

The fMRI underwent image re-orientation, motion and distortion correction, and nuisance signal regression (white matter, cerebrospinal fluid (CSF), and frame-wise displacement spikes) [18, 105, 43, 7]. Volumetric time series were mapped to the native FreeSurfer space using boundary-based registration [38, 23]. Vertex-level time series were averaged into parcels defined by the Schaefer atlas with 400 cortical regions [72]. Parcel-level time series were low-pass filtered to be less than 0.1 Hz. We computed the functional connectivity (FC) as the Pearson correlation matrix between parcel-level time series.

To estimate directed SC, we first ran our fingerprinting analyses (see below) to identify the optimal number of gene gradients. We fit subject-specific parameters in a hierarchical fashion. First, we fit a single model to the consensus SC and functional covariance to determine which sign each gene gradient should take. Since it is biologically implausible that gene gradients should operate with opposite polarity across brains in the same species, we fit subject-specific directionality models where gene gradient weights were constrained to have the same sign as the consensus result.

As an additional robustness analysis, we ran our model on an external public dataset, the MICA-MICs dataset [69]. This high-quality dataset uses the same processing pipeline we implemented on the HCP data, helping to ensure comparability between data processing.

### HCP Fingerprinting Analysis

We used test-retest fingerprinting analysis to optimize the number of gene maps for directionality prediction. That is, we sought the fewest gene maps that produced an asymmetry metric (*G***w**) with the best overall test-retest performance across three metrics: Intraclass Correlation Coefficient (ICC), Discriminability, and Top-1 Identification [22, 15]. For this purpose, we used two complementary approaches with the HCP dataset. First, we evaluated these metrics on *n* = 34 subjects from the HCP Test-Retest dataset; both test and retest have subject-specific SCs and 1 hour of scan time with the separation between scans being approximately 4 months. Second, we evaluated these metrics by splitting the fMRI time series into two ≈ 30-minute blocks using the full HCP dataset in subjects with the full 1 hour of scan time (*n* = 724).

We estimated feature-wise ICC (one-way, single-measures): for each feature, the *N* ×2 subject-by-session matrix was input to a variance-components ICC:

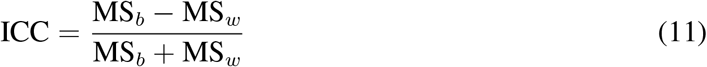

Here, MS*_b_* and MS*_w_* are the between-subjects and within-subjects mean squares, respectively. Uncertainty was quantified by nonparametric bootstrap resampling of subjects (1000 samples, with replacement) and was calculated as the 95% confidence interval for each feature’s ICC.

Second, we quantified multivariate discriminability between test and retest datasets. For each subject, we formed the Euclidean distance between that subject’s feature vector in the test (session-1) and retest (session-2), which corresponds to the “same-subject” distance, and compared it to the distances between that session-1 vector and all other subjects’ session-2 vectors, which correspond to the “different-subject” distances. Discriminability is defined as the proportion of pairwise comparisons in which the same-subject distance was smaller than the different-subject distance (i.e., the probability that a subject is closer to themselves than others across the two sessions). Statistical significance was assessed by a label-permutation test that randomly permuted subject indices in session-2 (5000 permutations) to generate a null distribution of the discriminability statistic. The permutation p-value was obtained as the proportion of the null distribution that was greater than the observed discriminability across subjects.

Third, we quantified Top-1 identification accuracy following a connectome fingerprinting framework [22]. Let *Y* ^(1)^*, Y* ^(2)^ ∈ R*^N^*^×*P*^ denote the subject-by-feature matrices from session 1 and session 2, respectively. We computed the Pearson correlations between all subject pairs across sessions and, for each subject, identified the single best-matching subject in the opposite session as the one with the maximal correlation. Identification accuracy was computed in both directions, and the final Top-1 score was defined as the average proportion of correct identifications across the two directions. Statistical significance was assessed with a label permutation test (5000 permutations), in which subject labels in one session were randomly permuted to generate a null distribution of Top-1 accuracy. To estimate feature importance for identification, we additionally computed a Finn-style differential power for each feature, based on the extent to which within-subject cross-session products exceeded between-subject cross-session products after feature-wise z-scoring across subjects in each session [22].

### Human Gene Ontology Analysis

After settling on the number of gene gradients (*k*) to use in our analysis, we used the asymmetry metric obtained from fitting the consensus (group-averaged) structural connectome to the consensus functional connectome: *g*^∗^ = *G*^(*k*)^**w**. We then calculated the cosine similarity between each gene expression map *E_i_* ∈ R*^n^* and the identified asymmetry metric *g*^∗^ as 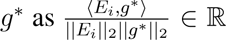. That is, for every gene, we obtained a measure of how well that gene is aligned (or anti-aligned) with the asymmetry metric. When the gene expression is aligned (anti-aligned) with *g*^∗^ we classified it as source-aligned (sink-aligned), since the expression is high (low) where the asymmetry metric is positive (negative).

To evaluate the distinct gene ontologies of source- and sink-aligned genes, we extracted the top 1,000 source- and sink-aligned genes, respectively, and entered them into the g:Profiler web-based interface [48]. We specified the gene universe “background” to be precisely the full list of genes within the Allen Human Brain Atlas (i.e., 15, 633 genes in our dataset). This provides a list of significant (*p <* 0.05, after FDR correction) gene ontology terms where the significance is based on a random sampling of our background. For interpretability, we limited our database search to the “GO” databases and the “Reactome” database. For each significant gene ontology term, g:Profiler provides an *intersection*, which is the number of genes from our empirical input gene set that were part of the term. Let the cardinality of the term intersection be denoted |*S*|.

Since brain maps tend to be spatially autocorrelated (SA), an additional statistical analysis is necessary on the g:Profiler results based on a SA-null distribution [17]. Therefore, we used a Moran spectral randomization to generate spatial-autocorrelation preserving null-distributions on the cortical maps with 1, 000 permutations [97, 99], and entered each of these maps into g:Profiler while exporting all results (including non-significant results) to compare against our empirical map results. Therefore, for each SA-null map, we have another measure of the gene ontology term intersection. Using these SA-null distributions, we can then calculate for each empirical term a SA-corrected p-value and a measure of SA-corrected enrichment:

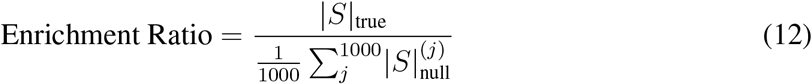

That is, the enrichment ratio calculates how many more genes, on average, appear in our empirical expression map compared to a SA-null-corrected background. In our results, we showed all terms that were significantly enriched relative to the background as determined by g:Profiler and we plotted their Enrichment Ratios compared to the SA-null distribution. Some terms were significant according to the generic background but only trending (0.05 *< p*_SA_ *<* 0.1) after calculating the SA-corrected p-value. We label these terms with a tilde (∼).

Since many terms are not independent of each other (i.e., both terms may describe closely related gene ontologies with nearly the same sets of genes), we grouped terms together based on a biological category after carefully evaluating each term’s biological relevance. This grouping should be considered an interpretive step and is not itself an outcome of the g:Profiler analysis.

### Network Source and Sink Analysis

To explore the source and sink organization of our estimated directed SC, we first assessed the skew connectivity, defined as *C̃*_skew_ = (C̃ _*C*^̃⊤^)/2. We used the rows of *C̃*_skew_ to visualize and qualitatively assess skew connectivity (i.e., the outgoing versus incoming structural relationships) within primary sensory areas and the thalamus. To summarize these skew edges across resting-state networks, we calculated *C̃*_(_*_R_*_)_ = *R*^⊤^*C̃*_skew_*R*, where *R* = [*r*_1_*, r*_2_*, …, r_m_*] given *r_i_* ∈ {0, 1}*^N^*^×1^.

Additionally, we investigated the overall degree imbalance of each node as the log ratio of out versus in degrees:

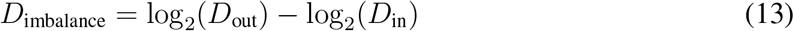

A positive degree imbalance indicates that the node has stronger outgoing connections than incoming connections, while a negative degree imbalance suggests the reverse. This measure captures the multiplicative imbalance between outgoing and incoming connectivity that can emphasize the sources or sinks that may have relatively few connections but are highly biased in their directionality.

### Angular Flow Analysis

Our directed SC and HONeD model imply a way to augment zero-lag functional covariance into a measure of functional “flow” through SC, which we term “angular flow” (AF). We compared AF against traditional directed/effective connectivity metrics in the Schaefer-100 atlas with 14 subcortical regions [72, 65]: namely lagged-FC, regression dynamic causal modeling (rDCM), and multivariate Granger causality (GC) [29, 10]. SC directionality for the 114-region atlas was identified for every subject using *k* = 5 gene gradients to keep the results homogeneous with the rest of the analyses. We calculated lagged-FC as the average sample (lagged) covariance across 10 lags. For rDCM and GC, we used established toolboxes [29, 10]. We statistically compared AF to these other directed functional connectivity metrics with both Pearson and Spearman correlations, reported together. When p-values were below computer precision, we denoted them as being *p* ≪ 0.001. When running comparisons, we used only the upper-triangular entries of the respective measure’s skew matrix.

When assessing resting-state connectivity, we evaluated the net flow for each node: Σ*_j_* Ω*_ij_* ^8^. Meanwhile, the resting-state functional connectivity principal gradient was calculated as the first non-trivial eigenvector of the Laplacian diffusion embedding in the same way as previous work [56]. We correlated these two maps using a Pearson correlation and evaluated its significance using a distance-dependent Moran spectral randomization on the cortical maps with 100, 000 permutations [97, 99].

We also assessed whether this alignment between AF and the principal FC gradient was greater when calculating AF using a directed SC compared to a symmetric (undirected) SC. We also sought to compare whether simple lagged-FC had net flow that significantly aligned with the same principal FC gradient across subjects. Alignment was calculated using a Pearson correlation. We compared each of these distributions with a paired t-test.

## Ethics Statement

Ethical approval was not required as confirmed by the license associated with the open access data.

## Acknowledgments

Data were provided by the Human Connectome Project, WU-Minn Consortium (Principal Investigators: David Van Essen and Kamil Ugurbil; 1U54MH091657) funded by the 16 NIH Institutes and Centers that support the NIH Blueprint for Neuroscience Research; and by the McDonnell Center for Systems Neuroscience at Washington University.

## Data and Code Availability

All data used in this study are publicly available from the sources cited in the Methods. Code is available at https://github.com/Raj-Lab-UCSF/DirectedSC.

## Author Contributions

BS: Conceptualization, Data Curation, Methodology, Formal Analysis, Software, Validation, Visualization, Writing – Original Draft, Writing – Review & Editing. FA: Methodology, Writing – Review & Editing. SN: Writing – Review & Editing. AR: Supervision, Funding Acquisition, Writing – Review & Editing.

## Supplemental Material

### Section 1: Extended Results

#### Statistics of Cross-Species Validation

To ensure results were not entirely driven by outlier edges, we also computed the Spearman’s *ρ* as a non-parametric correlation, which also showed robust positive correspondence between the model prediction and ground truth skew connectivity (*C. elegans*: *ρ* = 0.36; Mouse: *ρ* = 0.20; Macaque: *ρ* = 0.14; all *p* ≪ 0.001). We found similarly promising results when correlating the skew degree of empirical versus predicted networks (Pearson’s r — *C. elegans*: *r* = 0.67; Mouse: *r* = 0.60; Macaque: *r* = 0.54; Spearman’s *ρ* — *C. elegans*: *ρ* = 0.54; Mouse: *ρ* = 0.36; Macaque: *ρ* = 0.31; all *p* ≪ 0.001).

#### Identifying the optimal ***k*** in Cross-Species Validation

**Figure 6:**
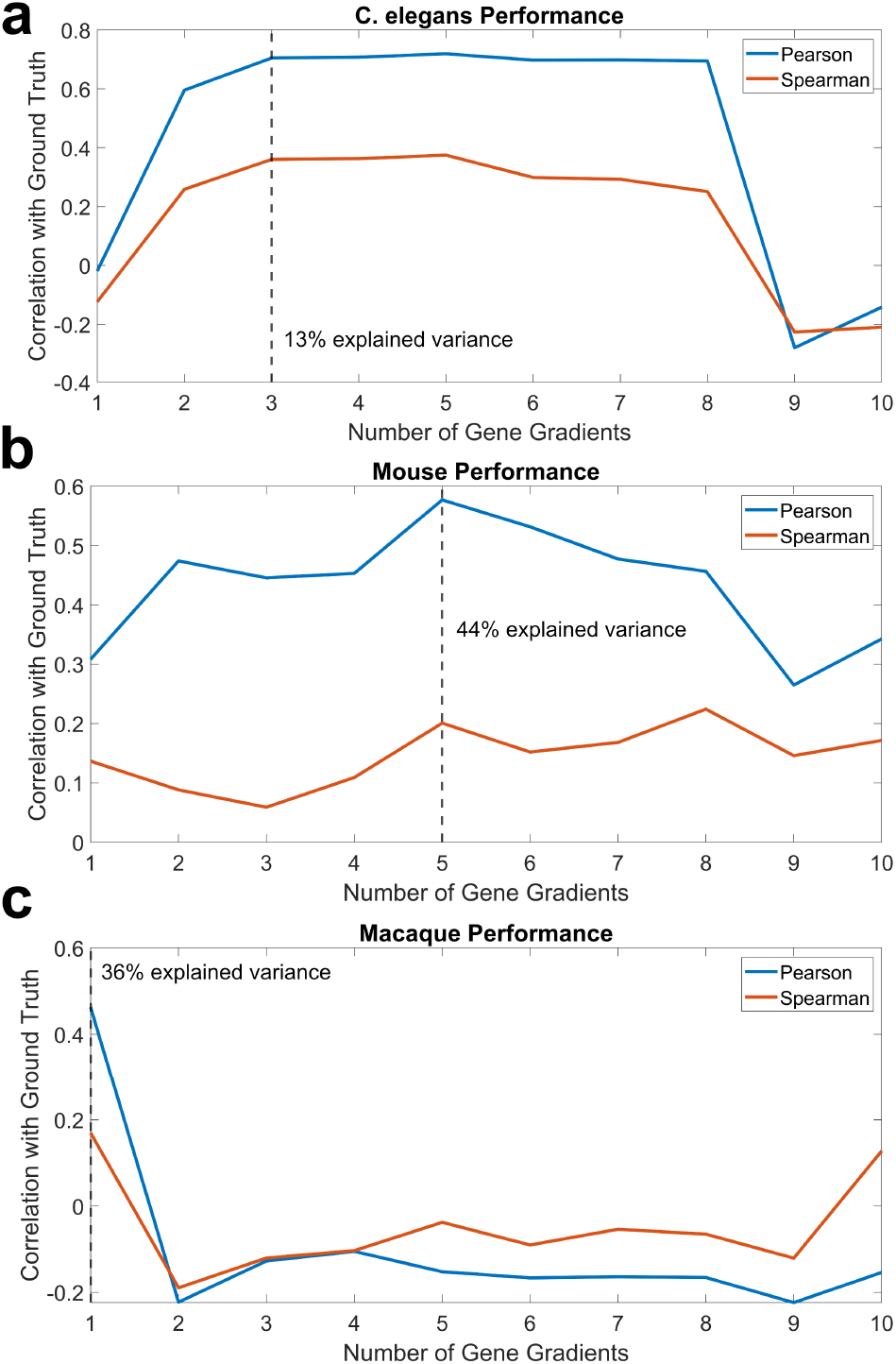
We evaluated the correlation between model-predicted and ground-truth directed (skew) connectivity with different numbers of gene gradients (*k*) ranging from 1 to 10. (**a**) *C. elegans* showed the best alignment with ground-truth at *k* = 3; (**b**) mouse showed the best alignment at *k* = 5; (**c**) macaque showed the best alignment at *k* = 1. The main results for the cross-species analysis are shown in Fig 2.

**Figure 7:**
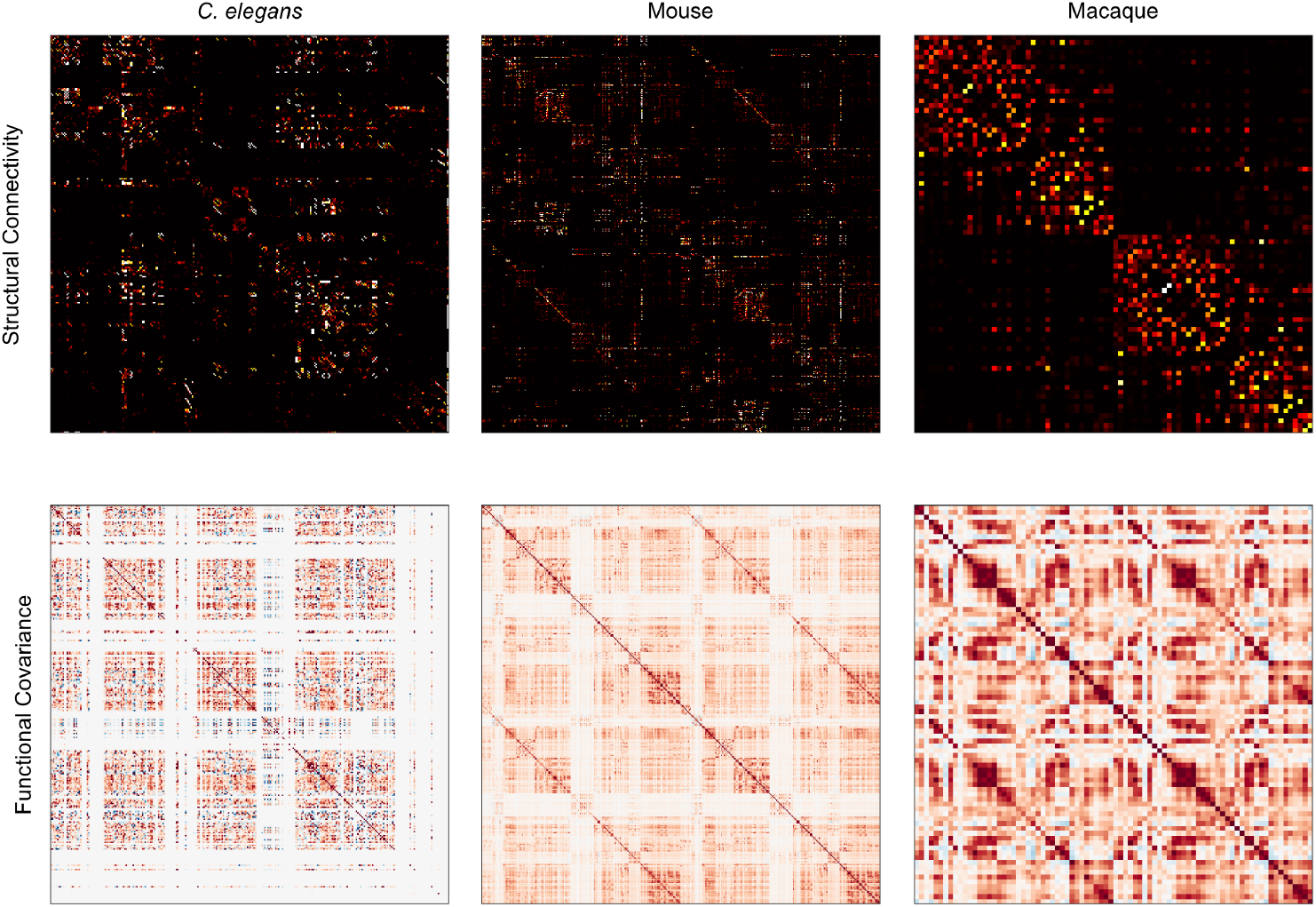
We visualize the structural connectivity and functional covariance matrices used to fit the directionality model for each species. The main results for the cross-species analysis are shown in Fig 2.

#### Human Fingerprinting and Replication

For the 34 test-retest dataset (Supplemental Fig. S 8a), *k* = 5 gene gradients showed a peak in ICC (0.46 ± 0.11), significant Discriminability (0.73 *p*_perm_ = 0), and significant Top-1 Identification (28%*, p*_perm_ *<* 0.001). For the 724 split fMRI subjects (Supplemental Fig. S 8b), we similarly found that *k* = 5 gene gradients had high ICC (0.51 ± 0.03), significant Discriminability (0.77*, p*_perm_ *<* 0.001), and a modest but significant Top-1 Identification (3%*, p*_perm_ *<* 0.001) (Supplemental Fig. S 8b). The most subject-specific identifiable regions in both datasets involved cingulate regions, left ventral prefrontal cortex, left inferior temporal gyri, and the hippocampus.

**Figure 8:**
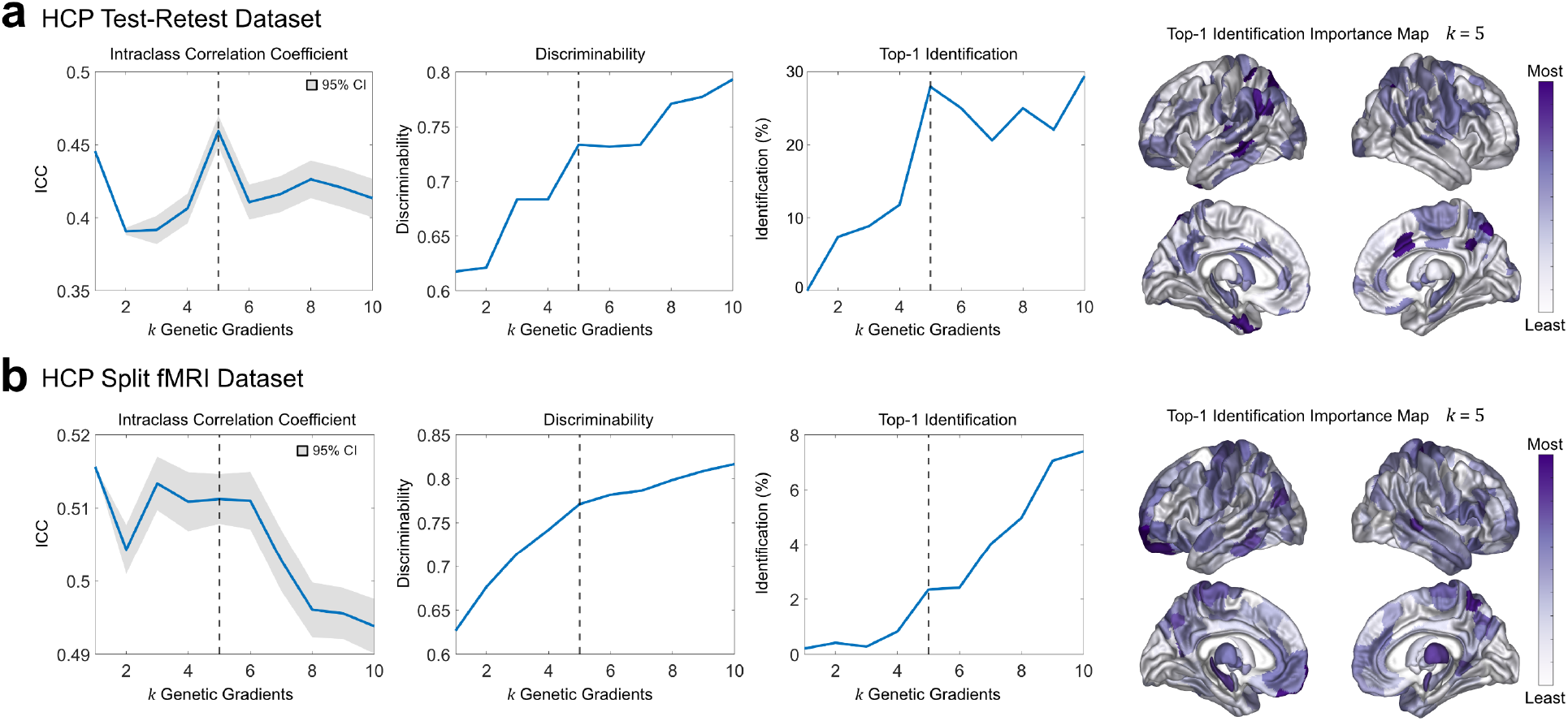
Human Asymmetry Test-Retest Fingerprinting. Fingerprinting on the fitted asymmetry metric for the (a) HCP Test-Retest dataset (*n* = 34), and (b) HCP split fMRI dataset (*n* = 724). We show (left to right) Intraclass Correlation Coefficient (ICC) with the 95% confidence interval shaded around the mean, the Discriminability score (all significant *p*_perm_ = 0), the Top-1 Identification metric (%), and the Top-1 Identification regional-importance map. The main results for the human analysis are shown in Fig 3.

**Figure 9:**
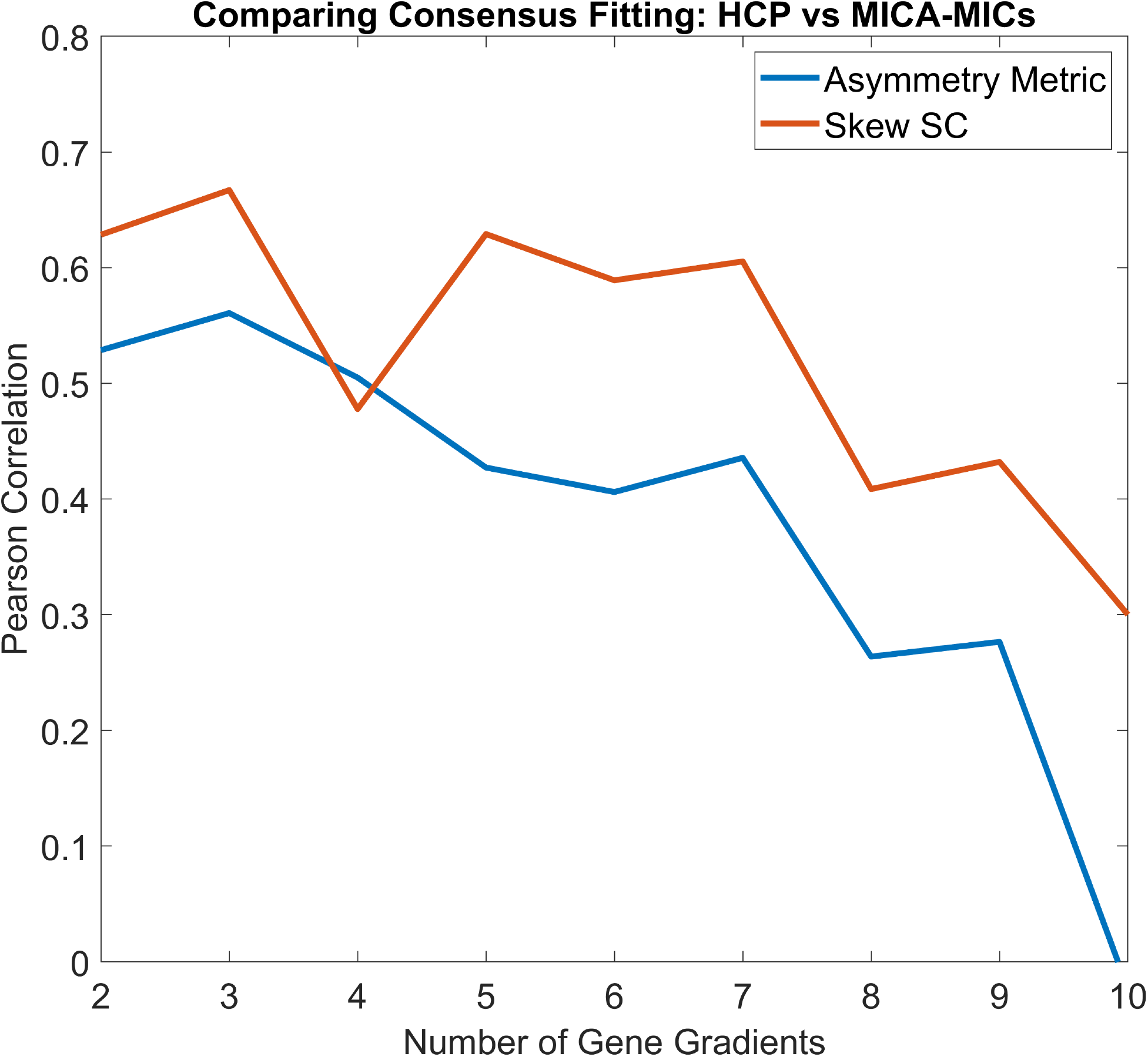
Human Directed SC Replication Dataset. We evaluated our model on the consensus connectome from the MICA-MICs dataset as an external validation. We evaluated the correlation between the predicted skew connectivity in the Schaefer-400 atlas for each dataset as well as the identified asymmetry metric. The main results for the human analysis are shown in Fig 3.

#### Top Sinks and Sources

**Table 1:** Top 10 regions with most and least directional imbalance from the group-averaged human connectome (Fig. 3e).

| Top 10 Sources |  | Top 10 Balanced |  | Top 10 Sinks |  |
| --- | --- | --- | --- | --- | --- |
| Name | Bias ($\log_2$) | Name | Bias ($\log_2$) | Name | Bias ($\log_2$) |
| Hippocampus (R) | 1.61 | Intraparietal Sulcus (R) | 0.0042 | Pallidum (R) | -2.16 |
| Posterior Cingulate (L) | 1.59 | Intraparietal Sulcus (L) | -0.0046 | Lateral Fusiform Gyrus (L) | -1.47 |
| Posterior Cingulate (L) | 1.47 | Dorsomedial Prefrontal Cortex (R) | -0.0049 | Intraparietal Sulcus (L) | -1.39 |
| Sup. Motor Area (L) | 1.44 | Postcentral Sulcus (R) | 0.0051 | Inferior Primary Motor Area (L) | -1.27 |
| Primary Motor Cortex (L) | 1.41 | Intraparietal Sulcus (L) | -0.0069 | Middle Frontal Gyrus (L) | -1.27 |
| Middle Frontal Gyrus (L) | 1.21 | Central Sulcus (L) | -0.0081 | Dorsal Primary Motor Cortex (L) | -1.25 |
| Primary Motor Cortex (L) | 1.11 | Middle Frontal Gyrus (R) | 0.0094 | Rostral Middle Frontal Gyrus | -1.21 |
| Medial Fusiform Gyrus (R) | 1.08 | Inferior Frontal Gyrus (R) | 0.012 | Secondary Auditory Area (L) | -1.20 |
| Posterior Cingulate (R) | 1.06 | Posterior Cingulate Sulcus (L) | 0.013 | Middle Frontal Gyrus (L) | -1.11 |
| Medial Fusiform Gyrus (L) | 1.05 | Caudate (L) | -0.015 | Pallidum (L) | -1.03 |

#### Extended Angular Flow Results

##### Statistical Comparison to Causal Discovery Methods

Lagged-FC significantly correlated with AF (*r* = 0.34*, p* ≪ 0.001, Spearman *ρ* = 0.34*, p* ≪ 0.001; Fig. 5b). We observed that GC had the strongest non-parametric relationship with AF (*r* = 0.22*, p* ≪ 0.001, Spearman *ρ* = 0.43*, p* ≪ 0.001; Fig. 5b) while rDCM showed more moderate but still significant correlations (*r* = 0.11*, p* ≪ 0.001, Spearman *ρ* = 0.13*, p* ≪ 0.001; Fig. 5b). Interestingly, GC and rDCM were negatively correlated by Pearson correlation but not by Spearman correlation, suggesting little monotonic relationship despite the Pearson association (*r* = −0.22*, p* ≪ 0.001, Spearman *ρ* = −0.0037*, p* = 0.77; Supplemental Fig. 10).

**Figure 10:**
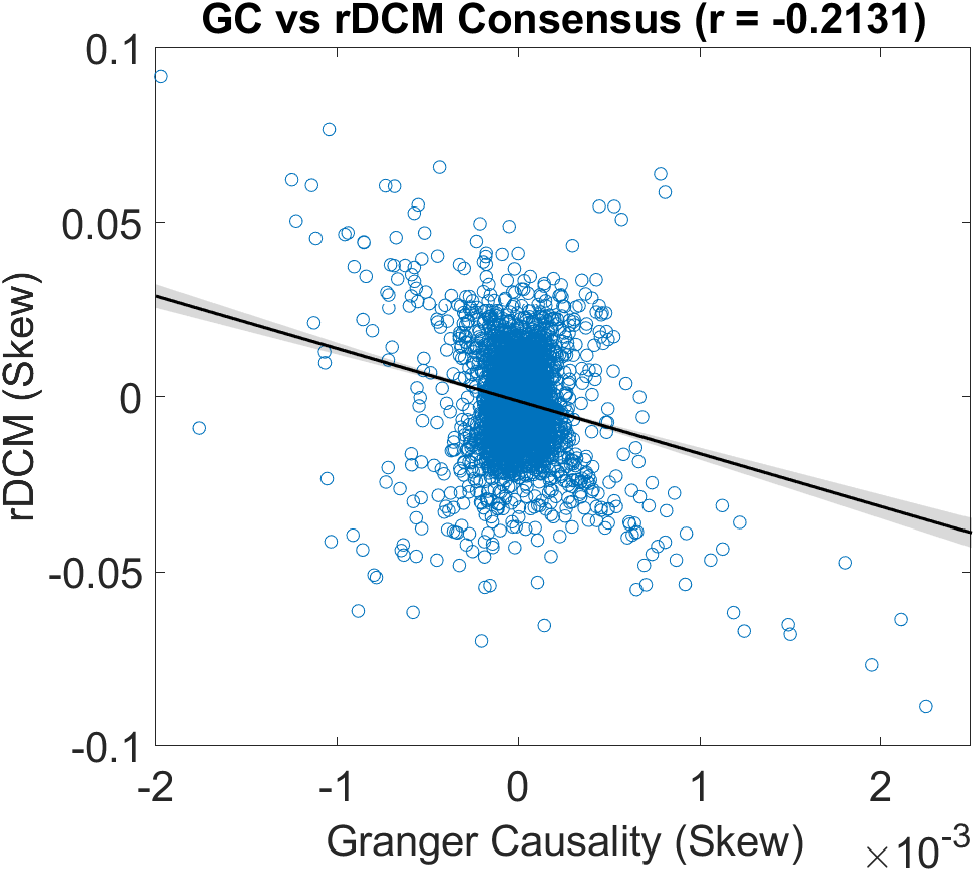
A comparison of the consensus rDCM skew-connectivity versus GC skew-connectivity. The main results for the AF analyses are shown in Fig 5.

#### Spin Test for AF Net-Flow and Principal Gradient

**Figure 11:**
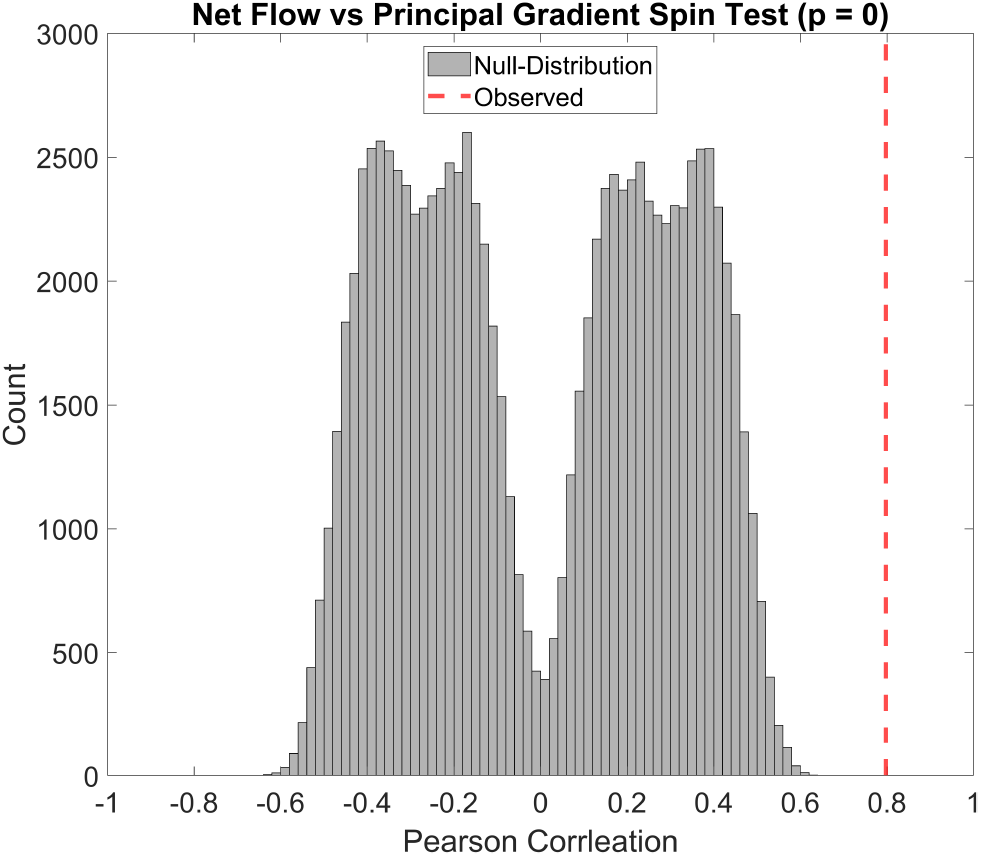
Moran spectral randomization results comparing the first principal gradient of undirected connectivity to the AF net-flow. The main results for the human AF analyses are shown in Fig 5.

#### Graph Laplacian Flow Operator

**Figure 12:**
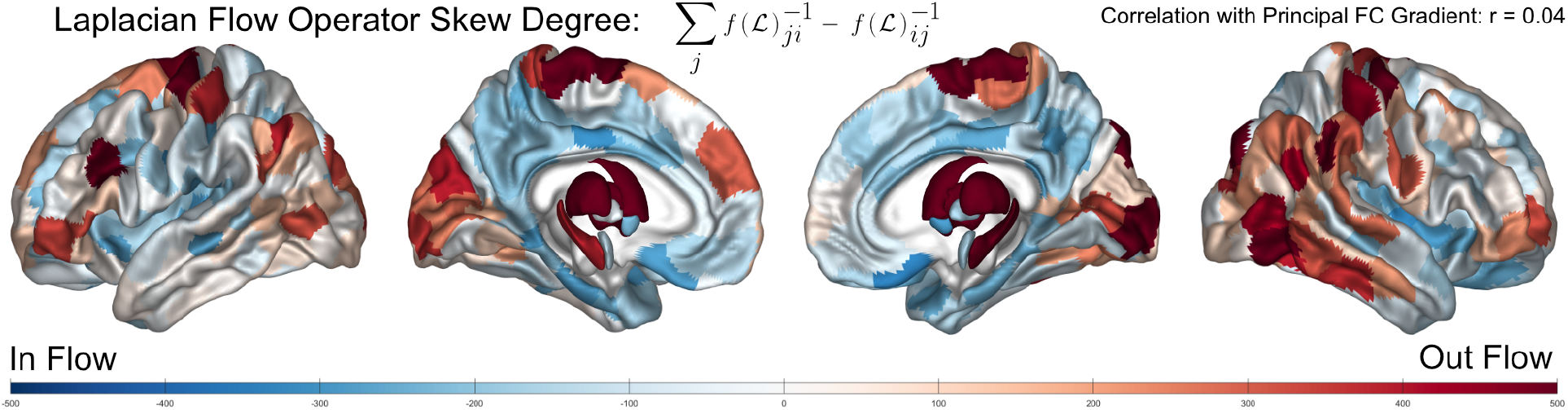
The net flow of the modeled diffusion operator in isolation. The main results for the AF analyses are shown in Fig 5.

#### Comparing Alignment with FC Gradient Across Cases

**Figure 13:**
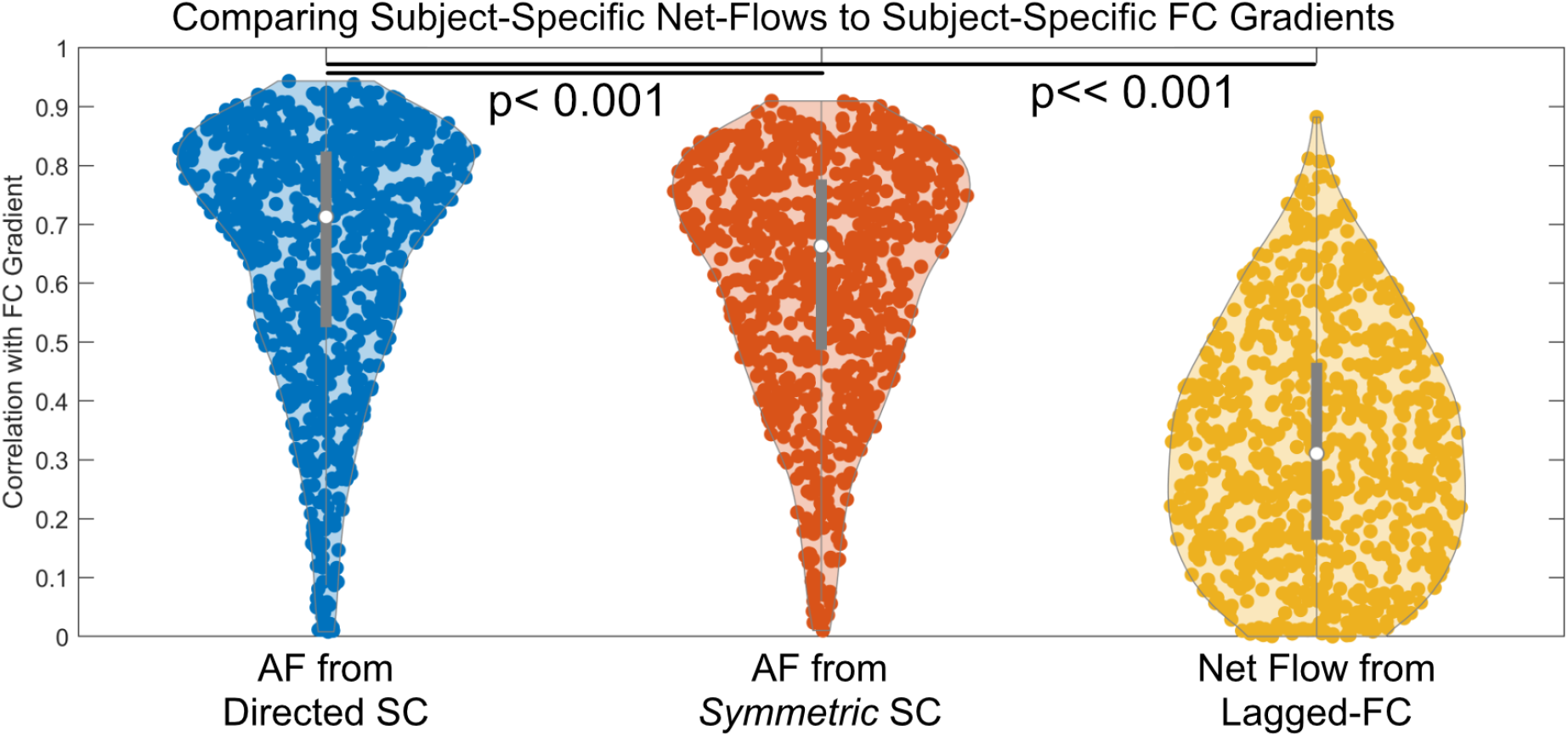
We statistically compared the predicted AF net-flow when calculated with the directed SC versus the original symmetric SC (*p <* 0.001, paired t-test), as well as against the net flow from the conventional lagged-FC. All comparisons were performed in the Schaefer 400 atlas. The main results for the AF analyses are shown in Fig 5.

### Section 2: Derivation of the Lyapunov Objective for Predicting Structural Directionality

#### Linear stochastic dynamics

We begin with a continuous-time linear stochastic model for the evolution of brain activity

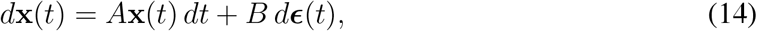

where **x**(*t*) ∈ R*^n^*is the vector of regional brain activity, *A* ∈ R*^n^*^×^*^n^* is the linear dynamics operator, *B* ∈ R*^n^*^×^*^r^* maps stochastic inputs into the observed state space, and *d**ɛ***(*t*) is a vector of Wiener-process increments satisfying

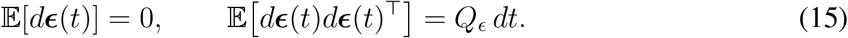

In our case, the deterministic dynamics are specified by the HONeD operator, so that

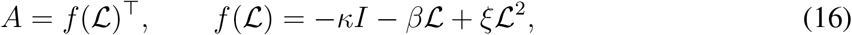

where L is the directed graph Laplacian and *θ* = {*κ, β, ξ*} denotes the diffusion parameters. The transpose appears because we write the state equation in the column-vector convention **ẋ** (*t*) = *f* (L)^⊤^**x**(*t*). For ease of notation, we abbreviate *f* (L; *θ*; **w**) ≡ *f* (L), where **w** denotes the parameters controlling structural directionality.

The effective covariance of the stochastic drive is

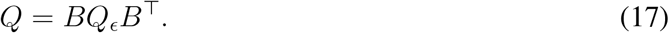

In the analyses below, we use an isotropic innovation model,

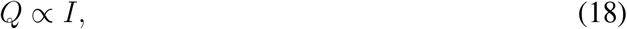

which assumes that unresolved local fluctuations inject variance into each region without imposing a preferred large-scale covariance structure. This choice is useful because the goal is to test whether the directed structural operator itself can explain the observed covariance structure, rather than allowing a flexible noise covariance to absorb that structure.

#### From state dynamics to covariance dynamics

The empirical functional covariance is modeled as the expected second moment

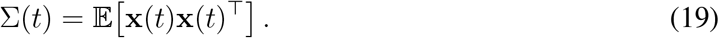

Since the data are mean-centered, this second moment is equivalent to the covariance. We now derive the differential equation governing Σ(*t*).

First, define the instantaneous outer product

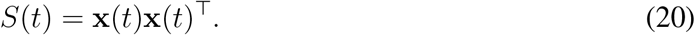

Using the stochastic product rule, also known as Itô’s product rule,

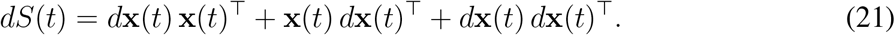

Substituting the stochastic dynamics from Eq. (14) gives

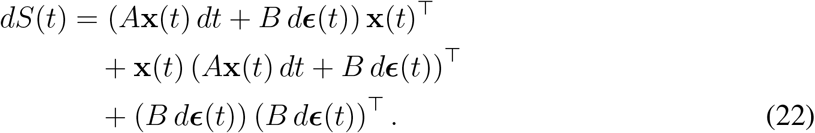

Expanding the first two terms,

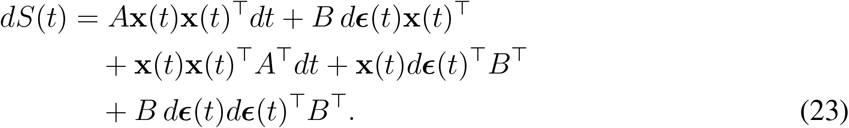

Taking expectations, the terms that are linear in *d**ɛ***(*t*) vanish because the Wiener increments have zero mean and are independent of the present state. Therefore,

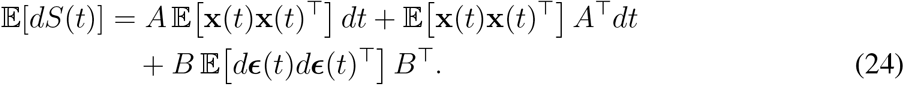

Using the definitions of Σ(*t*) and *Q* = *BQ_ɛ_B*^⊤^, we obtain

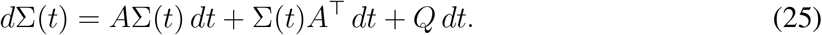

Dividing by *dt* gives the continuous-time differential Lyapunov equation

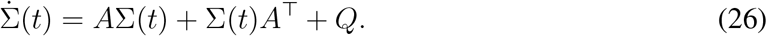

Substituting *A* = *f* (L)^⊤^, Eq. (26) becomes

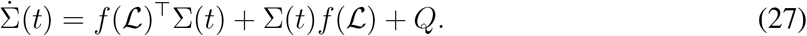

This is the Lyapunov equation used in our structural directionality objective.

#### Steady-state covariance constraint

If the linear dynamics are stable, meaning that the eigenvalues of *A* = *f* (L)^⊤^ have negative real part, then the covariance converges toward a stationary covariance Σ. At stationarity,

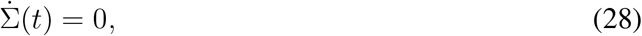

so Eq. (27) reduces to the algebraic Lyapunov equation

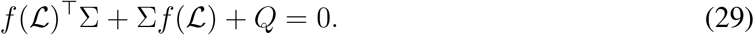

This equation expresses a balance between two contributions. The terms

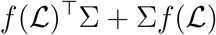

describe how the deterministic network dynamics dissipate, redistribute, or amplify covariance, whereas *Q* describes the covariance injected by stochastic innovations. At steady state, the observed covariance is one for which deterministic dissipation and stochastic injection are balanced.

## Footnotes

1 This parameterization is invariant under a constant shift. *Proof* : Consider the transformation *Â* → *Â* ≡ *sA* for any scalar *s >* 0. Then *ÂCÂ^−^*^1^ = (*sA*)*C*(*sA*)*^−^*^1^ = (*sA*)*C*(*s^−^*^1^*A^−^*^1^) = *ACA^−^*^1^. Equivalently in log coordinates, Â = diag(se^Gw^) = diag(e^(log s)+Gw^), and C̃_ij_ = C_ij_e^log s+(Gw)i−log s−(Gw)j^ = C_ij_e^(Gw)i−(Gw)j^.

2 The matrix power of a graph’s adjacency matrix, *G^n^* ∀*n* ∈ N, encodes walks of length *n* on the graph.

3 For ease of notation, we abbreviate the model *f* (L; *θ*; **w**) ≡ *f* (L). While we include the mathematical details with the actuation matrix (*B*) for completeness, the properties of *B* are neither explored in nor relevant to this work.

4 To keep our directionality convention homogeneous across networks, we used the transpose of the traditional dynamic causal model (DCM) framework where Σ(*τ*)*_ij_* = E[**x**(*t* + *τ*)**x**(*t*)*^⊤^*] implies a directed connection from *j* to *i*.

5 Note that in our HONeD model, *f* (L) is constrained to be Hurwitz.

6 The wedge symbol (∧) denotes the exterior product, also called the wedge product. Given two vectors, it produces an antisymmetric matrix that encodes the oriented area spanned by those vectors. In classical mechanics, angular momentum can be written as *L* = *r* ∧ *p*, where *r* is the position and *p* = *mṙ* is the momentum. In three dimensions, this bivector is equivalent via the Hodge-dual to the more familiar angular momentum vector *r* × *p*.

7 Data were provided by the Human Connectome Project, WU-Minn Consortium (Principal Investigators: David Van Essen and Kamil Ugurbil; 1U54MH091657) funded by the 16 NIH Institutes and Centers that support the NIH Blueprint for Neuroscience Research; and by the McDonnell Center for Systems Neuroscience at Washington University. Ethical approval was not required as confirmed by the license attached with the open access data.

8 Taking the sum of the AF along each row is equivalent to calculating the difference between the out-degree and in-degree of the flow connectivity *F* = −Σ*f* (L)*^−^*^1^.

